# Cadherin-11 Blockade Reduces Inflammation-driven Fibrotic Remodeling and Improves Outcomes After Myocardial Infarction

**DOI:** 10.1101/533000

**Authors:** Alison K. Schroer, Matthew R. Bersi, Cynthia R. Clark, Qinkun Zhang, Lehanna H. Sanders, Antonis K. Hatzopoulos, Thomas L. Force, Susan M. Majka, Hind Lal, W. David Merryman

## Abstract

**Background:** Over one million Americans experience myocardial infarction (MI) every year, and the resulting scar and subsequent cardiac fibrosis contribute to heart failure and death. A specialized cell-cell adhesion protein, cadherin-11 (CDH11), contributes to inflammation and fibrosis in rheumatoid arthritis, pulmonary fibrosis, and aortic valve calcification but has not yet been studied in the context of cardiac remodeling after MI. We hypothesized that targeting CDH11 function after MI would reduce inflammation-driven fibrotic remodeling and infarct expansion to improve functional outcomes in mice.

**Methods:** MI was induced by ligation of the left anterior descending artery in transgenic mice with reduced or ablated CDH11, wild type mice receiving bone marrow transplants from *Cdh11* transgenic animals, and wild type mice treated with a functional blocking antibody against CDH11 (SYN0012). Cardiac function was measured by echocardiography, expression of cell populations was quantified by flow cytometry, and tissue remodeling by altered histological assessment and transcription of inflammatory and pro-angiogenic genes by qPCR. Co-culture was used to assess interactions between cardiac fibroblasts and macrophages.

**Results:** MI increased transcription of *Cdh11* in non-cardiomyocyte cells. Mice with deletion of *Cdh11* and wild type mice receiving bone marrow transplants from *Cdh11* transgenic animals had improved cardiac function and dimensions after MI. Animals given SYN0012 had improved cardiac function, reduced tissue remodeling, and altered transcription of inflammatory and proangiogenic genes. Targeting CDH11 also reduced the number of bone marrow-derived myeloid cells and increased pro-angiogenic cells in the heart three days after MI, consistent with a decrease in transcription and expression of IL-6 in the infarct region. Cardiac fibroblast and macrophage interactions led to an increase in IL-6 secretion that was reduced with SYN0012 treatment in vitro.

**Conclusions:** Our findings suggest that CDH11-expressing cells contribute to inflammation-driven fibrotic remodeling after MI, and that targeting CDH11 with a blocking antibody improves cardiac function after MI. This improvement is likely mediated by altered recruitment of bone marrow-derived cells, thereby limiting the macrophage-induced expression of IL-6 by fibroblasts and promoting vascularization.

## Introduction

Over one million Americans experience a myocardial infarction (MI) every year, which significantly reduces cardiac function and potentiates the progression to heart failure by increasing the risk of recurrent infarctions.^1^ The process of infarct healing requires complex interactions between resident and recruited cells which must coordinate the clearance and replacement of damaged tissue with a stable and robust collagen scar to prevent cardiac rupture.

Immune cells – including monocytes and macrophages, among others – are recruited from the blood within the first few hours after MI and critically participate in the healing and remodeling process. Resident cardiac macrophages also participate in stabilizing the heart during cardiac disease.^2^ Following the successful clearance of necrotic tissue and cell debris by immune cells, resident mesenchymal cells – such as cardiac fibroblasts (CFs) – proliferate and differentiate into an activated, hyper-secretory, hyper-contractile, tissue remodeling phenotype known as myofibroblasts. This highly cellularized stage of cardiac remodeling, termed the granulation phase, is characterized by the resolution of inflammatory signaling and the transition to fibrotic remodeling and scar formation by activated myofibroblasts. In the following weeks, myofibroblasts deposit and remodel collagen into a compact scar, which can sustain the biomechanical integrity of the myocardial wall.^3^ However, excess inflammation and reparative activity can ultimately lead to expansion of the infarct area and further diminished cardiac function.

Development of treatment strategies for MI is made particularly challenging by the precise and necessary timing of both chemical signals and cellular activity. Many treatments targeting specific growth factor cascades have failed to maintain the delicate balance between necessary and excessive inflammation and fibrosis and often have adverse side effects on the surviving cardiomyocytes (CMs), causing additional loss of contractile potential. ^4–6^

Cadherin-11 (CDH11) is a cell-cell adhesion protein expressed by inflammatory cells and activated fibroblast-like cells in multiple inflammatory and fibrotic disease models – including rheumatoid arthritis, pulmonary fibrosis, and aortic valve calcification^7–9^ – but its function in infarct healing has not yet been studied. CDH11 engagement promotes the expression of the proinflammatory cytokine interleukin-6 (IL-6) as well as profibrotic signaling factors and myofibroblast markers, such as transforming growth factor beta 1 (TGF-β1), in diseased joints, lungs, and heart valves.^7–11^ CDH11 (or OB-cadherin) was originally described in osteoblasts and has been shown to affect cell migration and exfiltration in cancer studies,^12,13^ but the role of CDH11 in bone marrow-derived cell (BMDC) recruitment to the heart has not been studied. Thus, we hypothesized that genetic and pharmacologic targeting of CDH11 after MI would reduce inflammation-driven fibrotic scar expansion and improve cardiac outcomes.

## Methods

Detailed methods can be found in the Supplemental Materials. Briefly, MI was induced in mice by permanent coronary artery ligation.^14^ Cardiac remodeling post-MI was compared between *Cdh11* transgenic littermates (*Cdh11*^*+/+*^, *Cdh11*^*+/-*^, and *Cdh11*^*-/-*^), WT (C57BL6/J) mice receiving bone marrow from *Cdh11* transgenic donors, and between WT mice treated with either 10 mg/kg of a functional blocking antibody against CDH11 (SYN0012; used with permission from Roche) or an isotype control antibody (IgG2a). Antibody treatment was administered every four days, beginning one day after surgery, with the last treatment on day 17 after infarct.

Cardiac function was monitored by echocardiography, infarct stiffness was quantified by atomic force microscopy (AFM) and scar morphology (i.e., size and thickness) was quantified via a custom image processing analysis. Changes in CDH11-dependent cardiac gene transcription and protein localization throughout the heart was measured by quantitative polymerase chain reaction (qPCR) and immunohistochemistry, respectively. Flow cytometry was used to identify and quantify distributions of non-CM cell populations expressing CDH11 in the heart and peripheral blood. Isolated CFs and macrophages (MΦs) from WT mice were co-cultured to measure transcription and translation of inflammatory markers as measured by qPCR and enzyme linked immunosorbent assay (ELISA). Student t-tests and multi-way ANOVAs were used to determine statistical differences with a value of p < 0.05 considered significant.

## Results

To establish the therapeutic potential of targeting CDH11 in the heart after myocardial infarction, we first wanted to identify the cardiac cell populations on which CDH11 is expressed. Flow cytometric analysis of non-CM cardiac cell populations – including cardiac endothelial cells (CECs: CD45^−^CD31^+^), cardiac mesenchymal cells (CMCs: CD45^−^CD31^−^), and bone marrow-derived cells (BMDCs: CD45^+^) (**Figure S1** and **Figure S2**) – revealed that BMDCs constitute the majority of non-CM cells in the heart three days after infarct (86.3%), though by seven days the largest cardiac cell population is CMCs (52.3%), which is likely made up of a majority of activated myofibroblasts (**Figure 1A-B** and **Figure S3A**). Consistent with increased *Cdh11* transcription by non-CM cells after infarct (nearly 10-fold higher by seven days; **Figure S4A-B**), flow cytometry further revealed that each non-CM cell population had some level of CDH11 expression after MI and that the overall percentage of CDH11^+^ cells was increased by seven days (17.8% vs. 6.4% in Sham; **Figure S4C**). In particular, BMDCs constituted a significant percentage of CDH11^+^ cells in the heart three days after MI (65.3% vs. 30.5% in Sham), whereas CMCs made up the majority of CDH11^+^ cells at seven days (58.4%). Though the percentage of CDH11^+^ BMDCs was reduced back to Sham levels by seven days post-MI, the overall percentage of both CDH11^+^ BMDCs and CDH11^+^ CMCs were increased from day three and were higher than Sham (**Figure 1C**).

**Figure 1.**
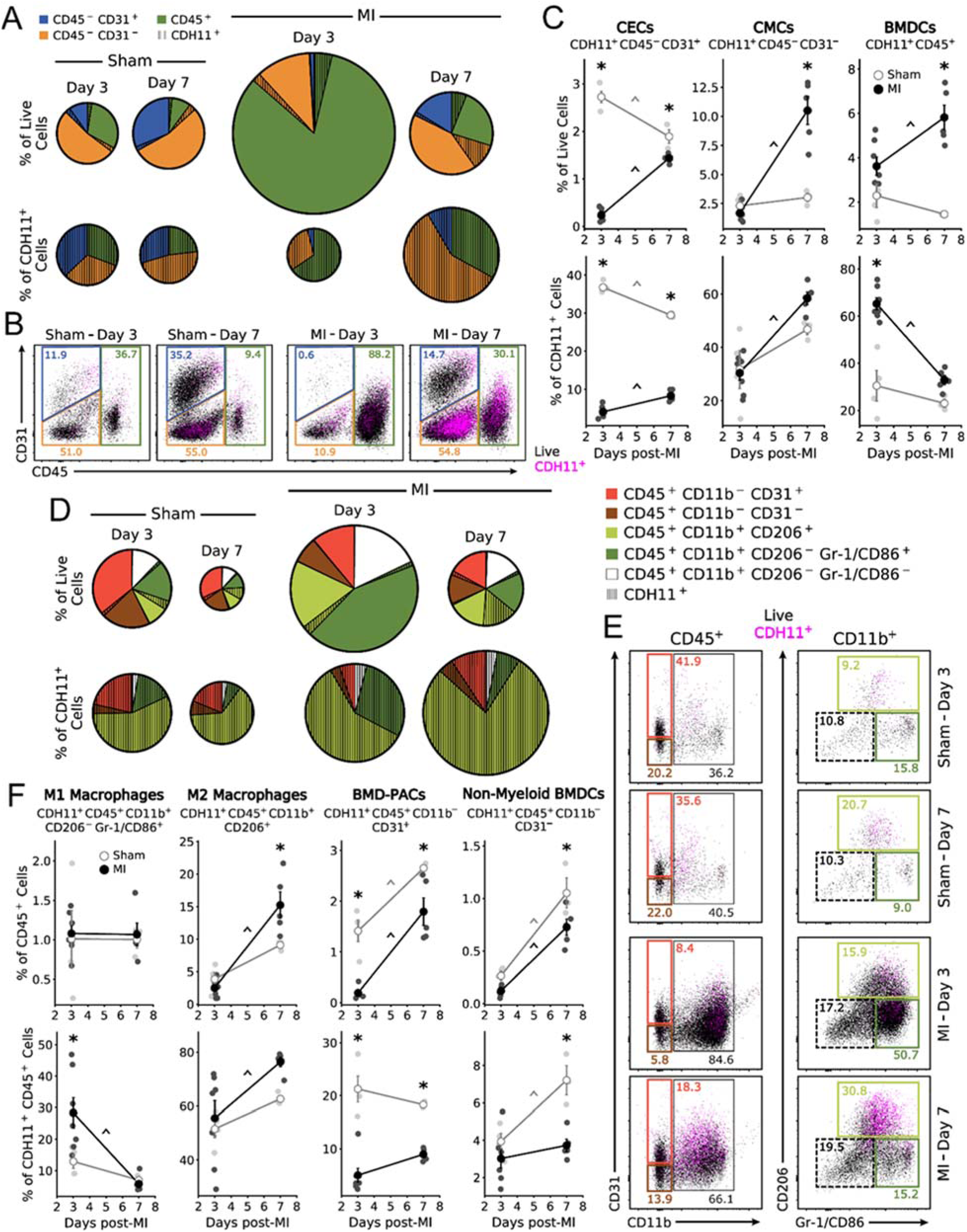
Specific cell populations in the heart express CDH11 post-MI. Non-cardiomyocyte cell populations – cardiac endothelial cells (CECs), cardiac mesenchymal cells (CMCs), and bone marrow derived cells (BMDCs) – express a baseline amount of CDH11 (hatched wedges) that is increased post-MI. Pie charts (**A**) are scaled by either total number of live single cells (top row) or total number of CDH11 expressing cells (bottom row), relative to Sham at day three. Representative dot plots (**B**) show changes in CDH11 expression (magenta) within each population (colored gates). CDH11 positive cells (**C**) within each population are shown as either a percentage of live cells or a percentage of all CDH11 positive cells. CDH11 expression in BMDC subpopulations (**D**) revealed predominant expression in M1- and M2-like macrophages (dark green and light green, respectively). Representative dot plots (**E**) show changes in CDH11 expression (magenta) within each subpopulation (colored gates). CDH11 positive cells (**F**) within each subpopulation are shown as either a percentage of all BMDCs or a percentage of all CDH11 positive BMDCs. * *p* < 0.05 between Sham and MI at the same time, ^ *p* < 0.05 over time; n = 3-7 per group.

We next analyzed our flow cytometry data to determine which BMDC subsets had CDH11 expression (**Figure 1D-F**). After infarct, we observed a significant increase in myeloid lineage cells (CD45^+^CD11b^+^) – including M1-like (CD45^+^CD11b^+^CD206^−^Gr-1/CD86^+^) and M2-like (CD45^+^CD11b^+^CD206^+^) macrophages – as well as a reduction in bone marrow-derived pro-angiogenic cells (BMD-PACs; CD45^+^CD11b^−^CD31^+^) and non-myeloid lineage BMDCs (CD45^+^CD11b^−^) – likely lymphocytes (**Figure S3A**). Though all BMDC subsets had detectable amounts of CDH11 expression, the largest fraction of CDH11 expressing BMDCs were myeloid lineage macrophages (**Figure 1D**). In particular, the percentage of CDH11^+^ BMDCs (i.e., CDH11^+^CD45^+^) that were M1-like macrophages was increased at day three, and the percentage of BMDCs that were CDH11^+^ M2-like macrophages was increased at day seven post-MI, relative to Sham (**Figure 1E-F**). After MI, the percentage of CDH11^+^ M2-like macrophages was increased from that at three days, and there was a significant reduction in the percentages of CDH11+ BMDCs that were BMD-PACs (at both three and seven days) and non-myeloid lineage cells (at seven days), relative to Sham (**Figure 1F**). A similar analysis of cells isolated from the peripheral blood post-MI confirmed that the vast majority (>90%) of cells were CD45^+^(**Figure S3B**), and revealed that only a small fraction (<2%) of cells expressed CDH11 (**Figure S4D**). Of the CDH11+ cells, little differences were observed between Sham and MI (**Figure S5**).

*Cdh11*^*+/+*^ mice had higher ejection fraction (EF) and LV mass as compared to *Cdh11*^*-/-*^ mice (**Figure 2A-B**), and both the LV mass and LV volume increased in *Cdh11*^*+/+*^ mice between day seven and day 21 after infarct but not in the *Cdh11*^*-/-*^ and *Cdh11*^*+/-*^ mice (**Figure 2B-D**). To further investigate the role of CDH11 in BMDCs, we induced MI in mice receiving bone marrow from *Cdh11* transgenic donors (**Figure 2E** and **Figure S6**). Though most of the mice (8 of 10) that received *Cdh11*^*-/-*^ bone marrow died before complete bone marrow reconstitution, *Cdh11*^*+/-*^ bone marrow recipients showed improved EF after 21 days and reduced LV mass after seven days post-MI, relative to *Cdh11*^*+/+*^ bone marrow recipients (**Figure 2F-G**). Left ventricular volume was not different between mice receiving either *Cdh11*^*+/+*^ or *Cdh11*^*+/-*^ bone marrow (**Figure 2H-I**).

**Figure 2:**
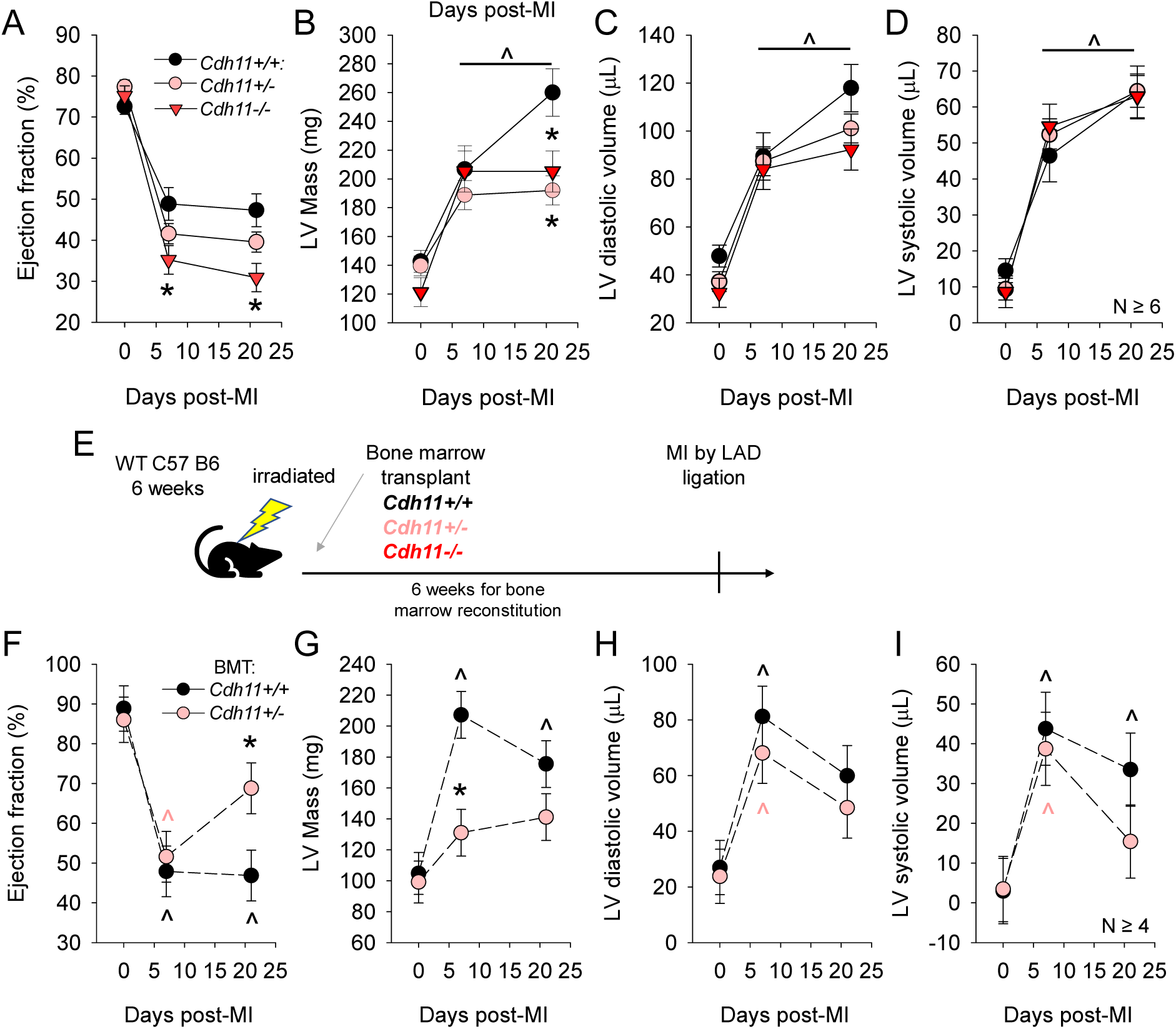
CDH11 mediates remodeling after MI in a partially BMDC-dependent manner. While ejection fraction (**A**), left ventricular (LV) mass (**B**), and LV volume (**C,D**) were significantly changed from baseline in all groups, *Cdh11*^*-/-*^ mice had significantly reduced ejection fraction despite a reduction in LV mass, as compared to *Cdh11*^*+/+*^. WT recipients of *Cdh11*^*+/-*^ bone marrow (**E**) had significantly improved EF at 21 days after MI (**F**), significantly lower LV mass at seven days after MI (**G**), and limited effect on LV volume (**H,I**) as compared to *Cdh11*^*+/+*^ bone marrow recipients. * *p* < 0.05 relative to *Cdh11*^*+/+*^, ^ *p* < 0.05 relative to previous time point (day 7 for **B-D** day 0 for **F-I**).

Having demonstrated a role for CDH11 in remodeling after MI, we hypothesized that blocking CDH11 adhesion and activity may therapeutically reduce MI-induced adverse ventricular remodeling and heart failure. Thus, MI was performed in mice treated with intraperitoneal injections of either a CDH11 blocking antibody (SYN0012) or IgG2a isotype control. EF (**Figure 3A**) and LV mass (**Figure 3B**) was preserved in SYN0012 treated animals relative to IgG2a for up to 56 days following MI. Further, the increased left ventricle dilation observed in the IgG2a treated mice was curtailed in the animals receiving SYN0012, resulting in preserved ventricular volume at both diastole (**Figure 3C**) and systole (**Figure 3D**) for up to 56 days post-MI. Though post-MI survival was improved with CDH11 blockade (82% for SYN0012 vs. 65% for IgG2a), it was not significant (p=0.16; **Figure S7**).

**Figure 3:**
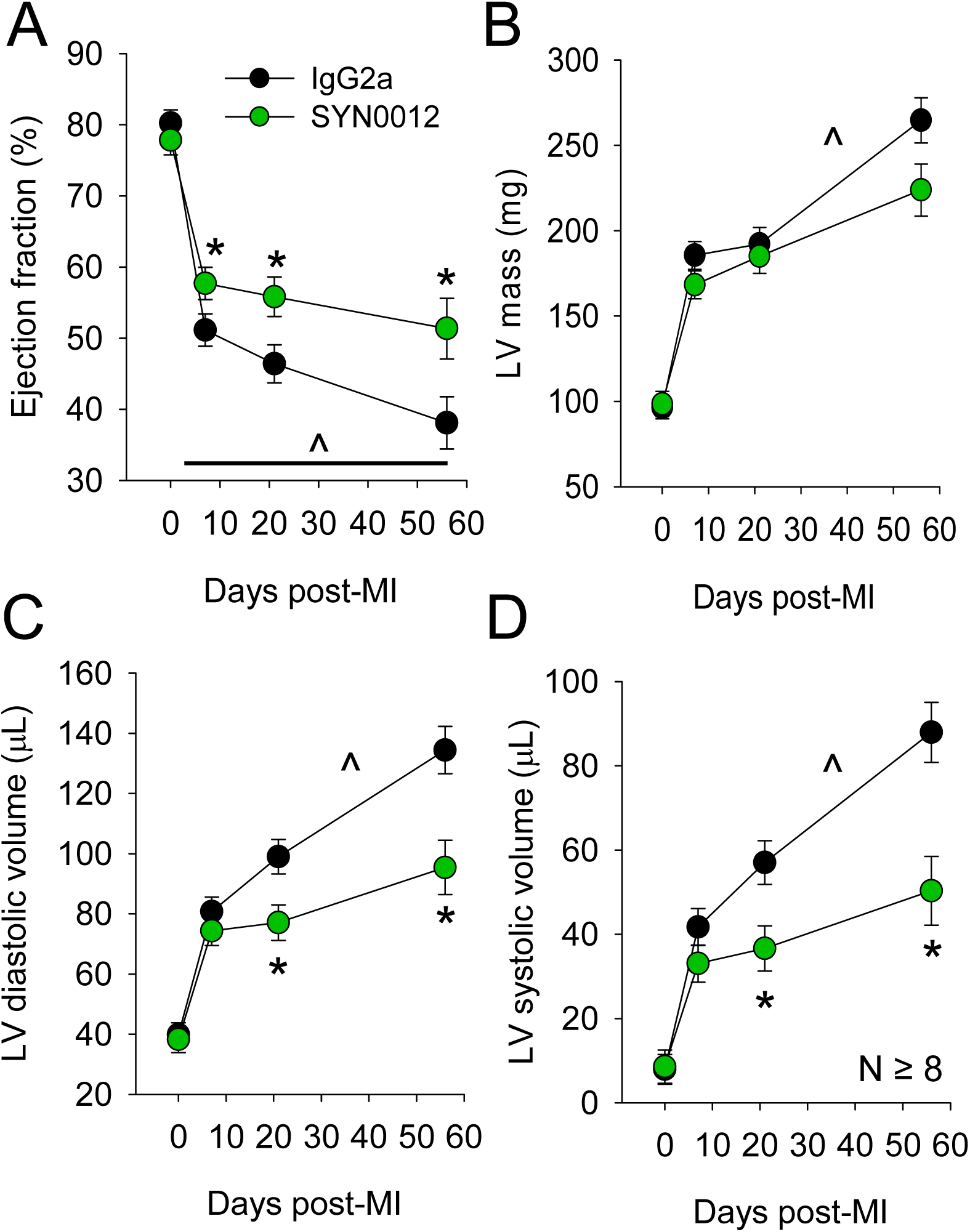
CDH11 blockade improves functional outcomes after MI. While ejection fraction (**A**), left ventricular (LV) mass (**B**), and LV volume (**C,D**) was significantly changed from baseline in all groups, SYN0012 treated animals had significantly improved ejection fraction and reduced LV expansion compared to IgG2a treated controls. * *p* < 0.05 between treatments, ^ *p* < 0.05 relative to previous time point.

Given the changes in left ventricular structure and function observed by echocardiography, we next wanted to quantify tissue mechanical properties and the extent of cardiac remodeling after infarct. In particular, by seven days post-MI, average stiffness of infarcted areas, as measured by AFM (**Figure S8**), in both SYN0012 and IgG2a groups was decreased relative to Sham myocardial stiffness (**Figure 4A**). However, by 21 days, infarct stiffness from both treatment groups was increased relative to Sham, with the range and maximum stiffness values from IgG2a-treated infarcts exceeding that of time-matched SYN0012-treated infarcts. Indeed, by 56 days post-MI, the average stiffness of SYN0012 treated infarcts was reduced relative to IgG2a and had returned to the Sham stiffness range (**Figure 4A**). Quantitative histological assessment (**Figure S9**) further revealed that infarcts from SYN0012-treated animals were thicker, more uniform, and spanned less of the left ventricular circumference than IgG2a at both 21 and 56 days post-MI (**Figure 4B-E**). There were no measured differences in cardiac hypertrophy or remote interstitial fibrosis between treatments (**Figure S10**), and despite differences in infarct remodeling, we interestingly observed no difference in CF contractility following SYN0012 treatment *in vitro*. Whereas *Cdh11* knockout reduces gel contraction by CFs (**Figure S11A**), SYN0012 treatment did not affect CF contraction (**Figure S11B**), suggesting a potentially critical role for CDH11-activation on both activated myofibroblasts and specific immune cell populations in determining the extent of cardiac remodeling after infarct.

**Figure 4:**
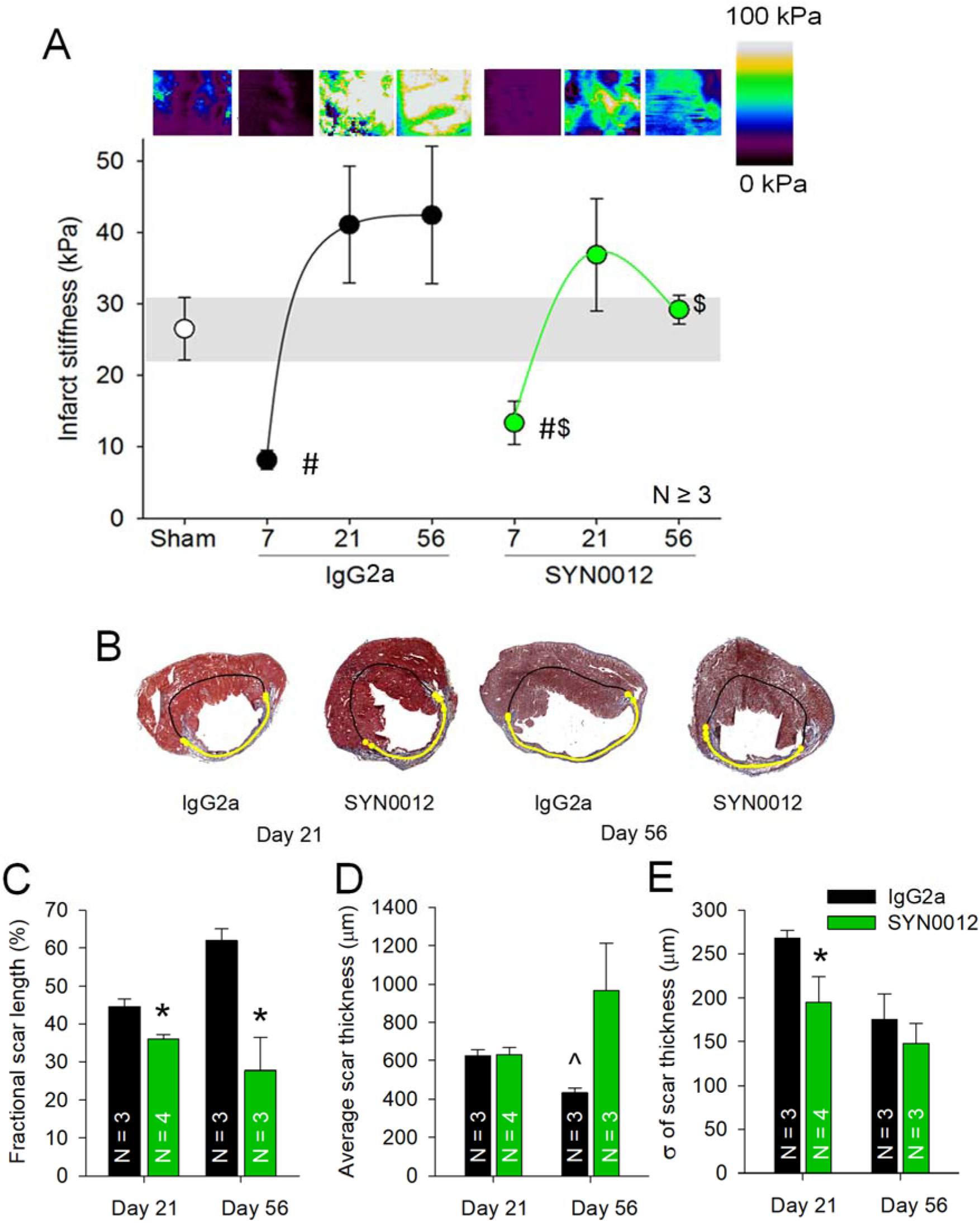
CDH11 blockade limits fibrotic remodeling after MI. Atomic force microscopy revealed a significant decrease in stiffness of the infarct relative to Sham myocardium at day seven after infarct (**A**). Representative stiffness color maps are depicted above the corresponding plot of average median stiffness. Stiffness variance was significantly reduced for IgG2a at seven days, but significantly increased at 56 days, relative to SYN0012 which was restored to the Sham range. Representative trichrome sections denote scar location (yellow line) (**B**). Images were used to quantify the infarct length (**C**), average thickness (**D**), and thickness variation, as measured by the standard deviation along the infarct length (**E**). * *p* < 0.05 between treatments, # *p* < 0.05 relative to Sham, ^ *p* < 0.05 relative to previous time point, $ *p* < 0.05 relative to variance of IgG2a.

To determine which cells contribute to the preserved cardiac remodeling in response to pharmacologic CDH11 blockade, we again used flow cytometry to evaluate the percentages of non-CM cells in the heart and peripheral blood after infarct (**Figure 5**). Though differences were observed between all populations at three and seven days, SYN0012 treatment did not alter the percentages of non-CM cardiac cell populations – CECs, CMCs, and BMDCs – relative to IgG2a, despite trends toward decreased CMCs and increased BMDCs after infarct (**Figure 5A-C**). Further examination of BMDC subsets revealed a decrease in myeloid lineage cells in response to SYN0012 treatment at three days post-MI, consistent with a reduction in both M1-like and M2-like macrophages (**Figure 5D-F**). Though not significant, the ratio of M1:M2 macrophages in the heart was reduced by 19.8% at day three (1.11 for IgG2a vs. 0.89 for SYN0012) and 33.3% at day seven (0.66 for IgG2a vs. 0.44 for SYN0012), suggesting SYN0012 treatment preferentially alters the balance of macrophage polarization in the heart toward a more pro-resolving inflammatory environment after infarct (**Figure 12A-C**). In addition, SYN0012 resulted in an increase in BMD-PACs and a non-myeloid BMDCs at day three; differences relative to IgG2a in all populations were gone by day seven. The distribution of circulating cells in the peripheral blood was largely unaffected by SYN0012 treatment, aside from more BMD-PACs in the peripheral blood at day seven after infarct, relative to IgG2a treatment (**Figure S13D-F**).

**Figure 5.**
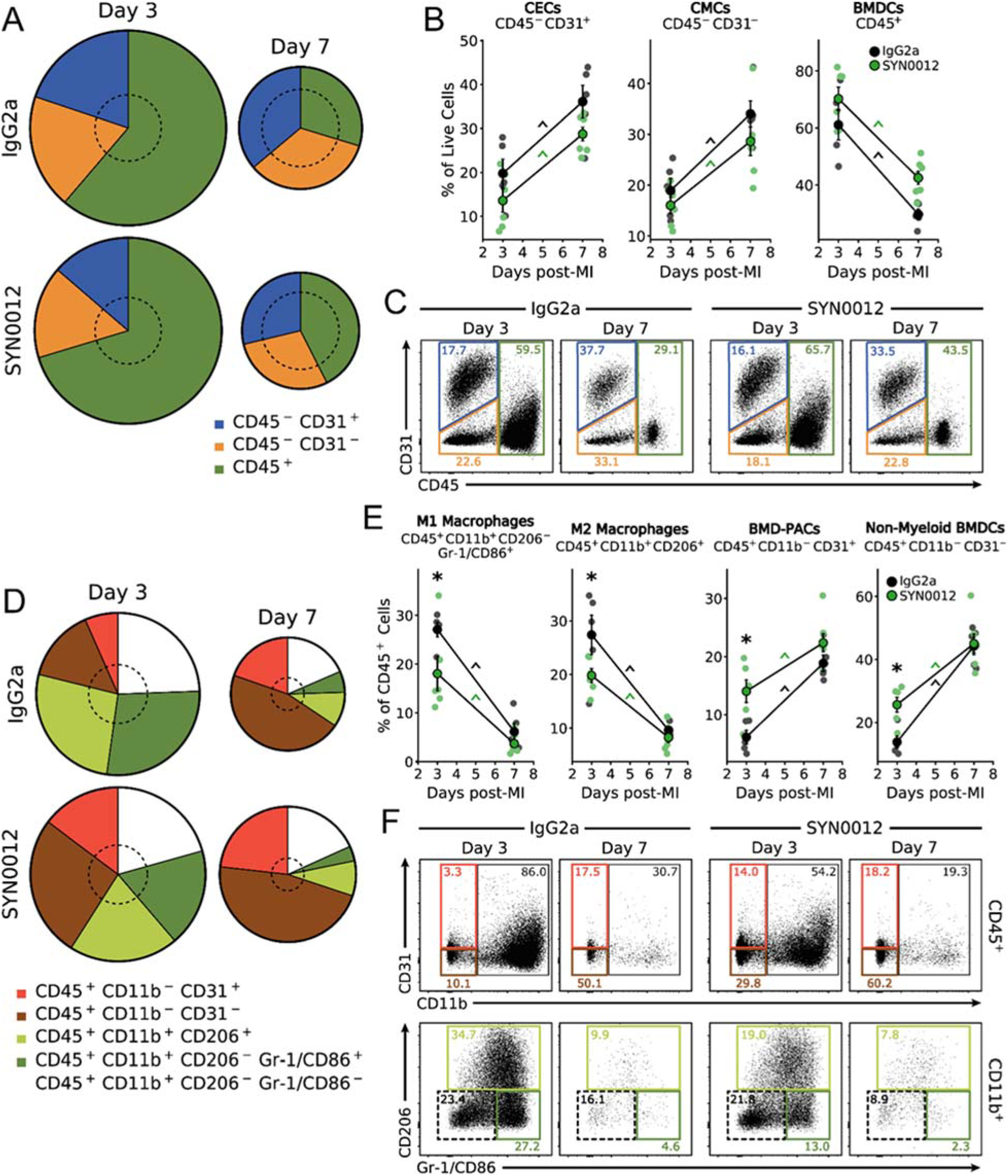
CDH11 blockade modulates expression of specific cell populations post-MI. SYN0012 treatment does not significantly alter the percentages of non-cardiomyocyte cell populations (CECs, CMCs, and BMDCs) in the heart post-MI, relative to IgG2a isotype control (**A-B**). Pie charts are scaled by the number of live single cells for each treatment and time; dotted circles denote the relative number of cells in Sham hearts. Representative dot plots (**C**) show changes in expression of each population (colored gates). Separation of BMDC populations (**D**) revealed that SYN0012 results in a significant reduction in M1- and M2-like macrophages in addition to increased BMD-PACs and a non-myeloid BMDCs (likely lymphocytes) at day three after infarct; differences relative to IgG2a in all populations were gone by day seven (**E**). Representative dot plots (**F**) show changes in expression of each subpopulation (colored gates). * *p* < 0.05 between treatments at the same time, ^ *p* < 0.05 over time; n = 3-7 per group.

We next examined transcriptional levels of multiple inflammatory and fibrotic genes over the 21 day time course of remodeling after MI (**Figure 6** and **Figure S14**). SYN0012 treatments decreased transcription of the pro-inflammatory cytokine *Il6*, relative to IgG2a, at three days post-MI (**Figure 6A**), and immunostaining confirmed that there was a reduction in IL-6 expression in non-CM cells in the infarct region of SYN0012 treated hearts (**Figure 6B**). Transcription of *Mmp13*, a matrix-metalloproteinase responsible for breakdown of collagenous tissue, was restored to baseline levels by day seven in the SYN0012 treated animals, but persisted in the IgG2a treated animals, relative to Sham, until day 21 (**Figure 6C**). Interestingly, SYN0012 treatment reduced transcription of pro-angiogenic signaling factors fibroblast growth factor (*Fgf2)* and vascular endothelial growth factor a (*Vegfa1)*, relative to IgG2a. In particular, IgG2a treated hearts had increased *Fgf2* transcription, relative to Sham, at all measured time points, but was only increased at 21 days in SYN0012 treated hearts (**Figure 6D**). While *Vegfa1* transcription was not different from Sham at any time point, SYN0012 treatment resulted in a reduction at day three after infarct (**Figure 6E**); however, SYN0012 treated animals appeared to have more muscularized vessels – likely arterioles – in the infarct region at 21 days post-MI (**Figure 6F**). Smooth muscle α-actin (α-SMA) is a contractile form of actin expressed in both myofibroblasts and smooth muscle cells which surround arteries and arterioles.

**Figure 6:**
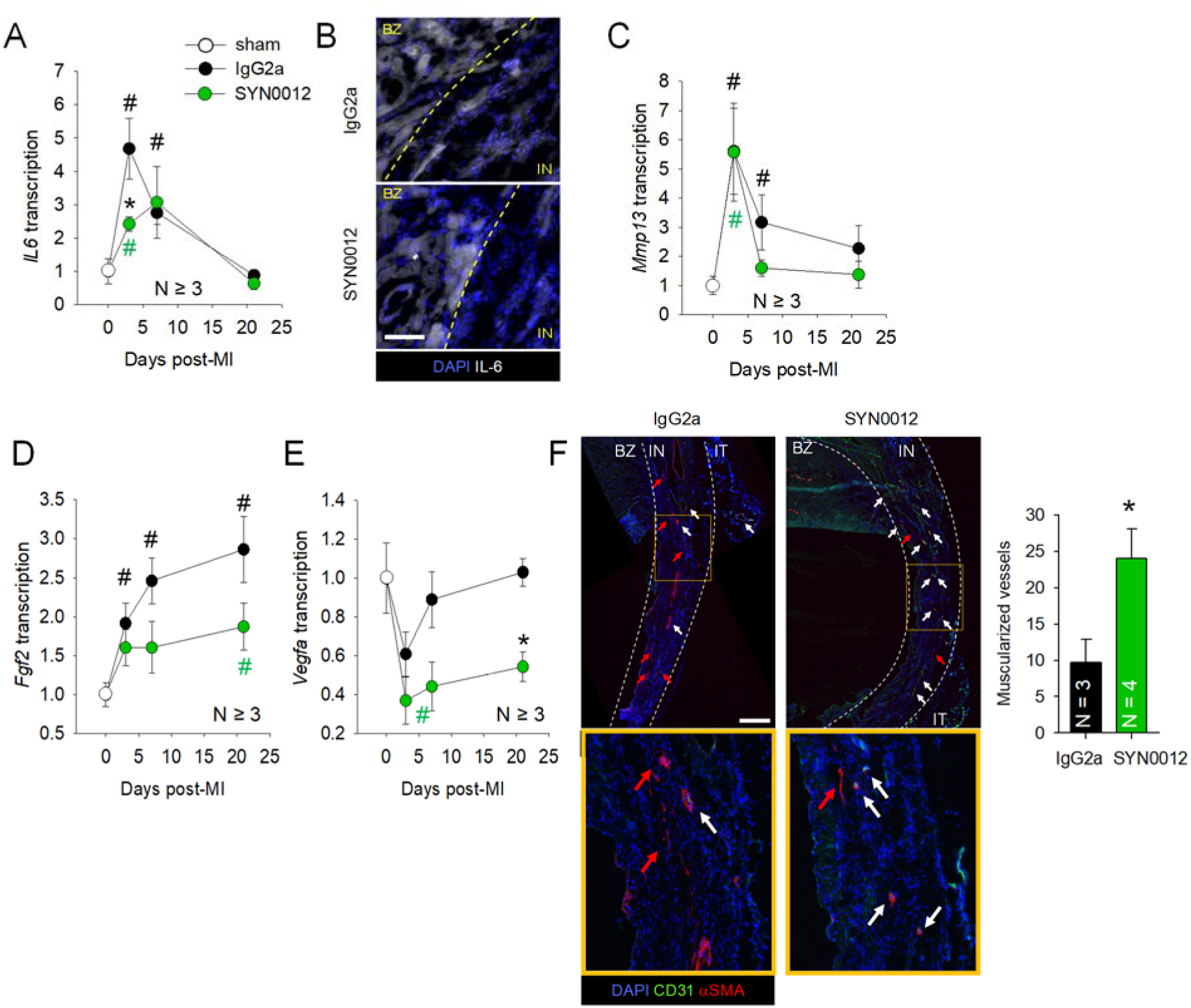
CDH11 blockade reduces inflammatory signaling and improves vascular maturity after MI. Transcription of *Il6* (**A**) was significantly decreased at three days post-MI following SYN0012 treatment, relative to IgG2a. Immunostaining of the border zone (BZ) and infarct (IN) confirmed decreased IL-6 expression three days after infarct, primarily in the non-CM cells of the infarct (**B**). Transcription of *Mmp13* (**C**) returned to baseline levels by day seven in SYN0012 treated hearts, whereas IgG2a returned to Sham by day 21. Transcription of both *Fgf2* (**D**), and *Vegfa1* (**E**) were all decreased by SYN0012, relative to IgG2a. Analysis of histological sections showed an increase in the number of muscularized found in the infarct region of SYN0012 treated animals (**F**). Red arrows indicate myofibroblasts, white arrows indicate arterioles, and the yellow box is a callout magnified below. Scale bar for B is 100 im and scale bar for F is 500 mm. * *p* < 0.05 between treatments, # *p* < 0.05 relative to Sham; color significance marker denotes group.

Based on our findings of increased CDH11 expression in BMDC macrophages (MΦ) and resident CMCs (which we expect to be primarily CFs), we next wanted to determine if interactions between these two cell populations may regulate the expression or transcription of pro-inflammation, pro-fibrotic, or pro-angiogenic factors (**Figure 7**). Indeed, *in vitro* CF-MΦ co-cultures of with varying CF:MΦ ratios confirmed previously reported findings that CDH11-dependent interactions between CFs and MΦs promote IL-6 secretion by CFs (**Figure 7A**).^15^ Of note, the MΦs alone expressed very low levels of IL-6. Treatment with SYN0012 reduced, but did not prevent, secretion of IL-6 by CFs (**Figure 7B**). qPCR further revealed that *Mmp13* transcription was increased in a CF:MΦ dependent manner, but was not affected by SYN0012 treatment (**Figure 7C-D**). Interestingly, *Tgfβ1* transcription was increased by CF-MΦ interactions and further increased with SYN0012 treatment (**Figure 7E**), whereas transcription of pro-angiogenic factors *Fgf2* and *Vegfa1* were not affected by varying CF: MΦ ratios (**Figure S15A-B**). *Arg1* transcription a marker of M2 macrophage polarization – was increased by treatment with SYN0012, though the transcript level of M1 and M2 polarization markers *Cd14* and *Mrc1* were not altered by SYN0012 treatment (**Figure S15C-D**).

**Figure 7:**
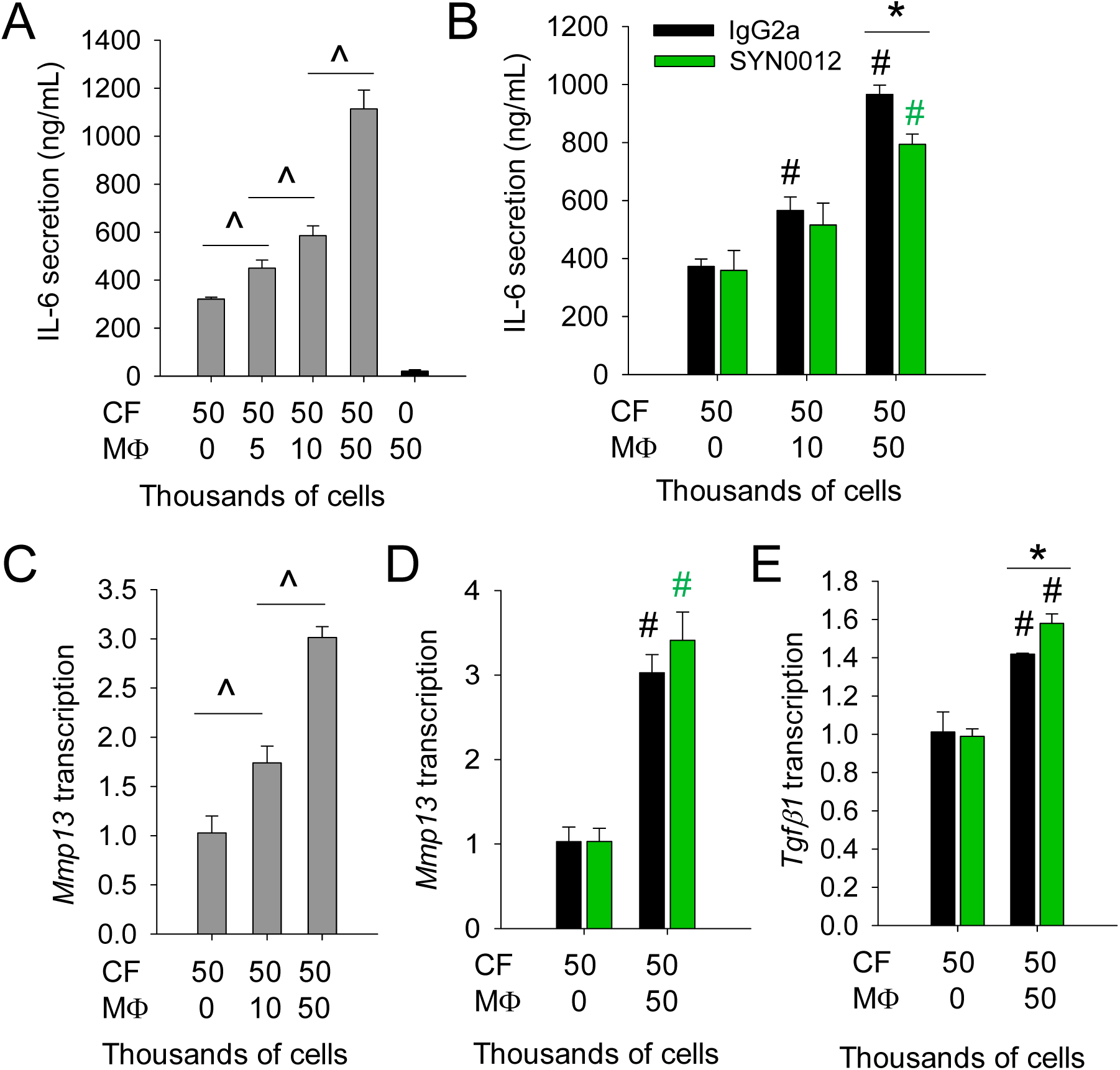
Co-culture of macrophages and cardiac fibroblasts increases proinflammatory and profibrotic signaling. Co-cultures of macrophages (MΦ) and cardiac fibroblasts (CF) promote secretion of IL-6 by CFs in a MΦ-dependent manner (**A**). CDH11 blockade by SYN0012 treatment significantly reduced IL-6 secretion in MΦ-CF co-cultures (**B**). Transcription of *Mmp13* in significantly increased by the addition of MΦ to CF culture (**C**) but was not significantly affected by SYN0012 treatment (**D**). TGF-β1 transcript was significantly increased in co-cultures and was further increased by SYN0012 treatment (**E**). N=3 for all samples, ^ *p* < 0.05 between MΦ conditions, # *p* < 0.05 relative to CF-only control, * *p* < 0.05 between treatments.

## Discussion

Our findings reveal that CDH11 is expressed in the ischemic heart, primarily in non-CM cells, and suggest a functional role in resolving tissue breakdown and promoting myocardial remodeling. *Cdh11* transcription is nearly 10-fold higher at seven days after MI, as compared to Sham (**Figure S4A-B**), suggesting a prominent role in activated myofibroblasts which are particularly active when the bulk of scar formation occurs between two and three weeks after MI.^3^ However, the observation that *Cdh11* is also significantly upregulated as early as three days after infarct suggests that other cell types – particularly those derived from infiltrating inflammatory cells, such as macrophages – may also contribute to CDH11-mediated cardiac remodeling. Flow cytometric analysis confirmed that CDH11 is expressed in the ischemic heart and further revealed that there is baseline CDH11 expression in approximately 5% of all non-CM cells (**Figure S4C**) distributed evenly between CECs, CMCs, and BMDCs. Three days after MI, the majority of CDH11^+^ cells were BMDCs whereas by day seven the majority of CDH11^+^ cells were CMCs, consistent with the time course of inflammation resolution and myofibroblast activation (**Figure 1A-C**). Note that increased CDH11 expression is known to be a hallmark of myofibroblast phenotype in other cardiovascular cell types, such as aortic valve interstitial cells.^16^ At both three and seven days post-MI, the majority of CDH11^+^ BMDCs were myeloid lineage macrophages with either M1-like or M2-like polarization, with the highest expression in M2-like macrophages (**Figure 1D-F**). The time-dependent expression of CDH11 in the ischemic heart suggests that targeting of CDH11 after MI has the potential to affect both early tissue breakdown and inflammation – driven by macrophages – as well as later collagen deposition and remodeling – driven by myofibroblasts to result in significant functional improvement after infarct.

*Cdh11*^*-/-*^ transgenic mice had reduced cardiac function after MI, which suggests an important role for CDH11 in myocardial maintenance and the response to MI (**Figure 2A**). *Cdh11*^*-/-*^ animals have been shown to have reduced ECM expression and tissue integrity, which likely contributes to this effect.^17^ While LV mass and volume continue to increase in *Cdh11*^*+/+*^ mice between day seven and 21, the LV dimensions of the *Cdh11* transgenic animals do not change significantly over this period, suggesting that CDH11 contributes to the fibrotic remodeling phase of MI healing (**Figure 2B-D**). Transplantation of bone marrow from *Cdh11*^*-/-*^ donor was fatal to most of the WT recipients, likely due to impaired localization of hematopoietic cells to the bone marrow (**Figure S6**).^12^ However, surviving *Cdh11*^*+/-*^ bone marrow recipients had improved EF three weeks after MI, relative to *Cdh11*^*+/+*^ bone marrow recipients (**Figure 2F-I**). Overall, these data suggest that CDH11 has important functional roles in multiple resident and recruited cell types, particularly in localization of BMDCs and activity of myofibroblasts. However, permanent loss of CDH11 in this mouse model may cause compensatory changes that increase the severity of cardiac damage after MI. *Cdh11*^*-/-*^ valve interstitial cells have reduced contractility,^16^ which we also observed in *Cdh11*^*-/-*^ CFs, but notably did not observe in wild type CFs treated acutely with SYN0012 (cf. **Figure S11**).

Previous studies have shown that genetic and pharmacologic targeting of CDH11 improves functional responses in both fibrotic lung disease and aortic valve calcification.^8,9^ Herein, CDH11 blockade preserves cardiac function as early as seven days after MI and prevents the continued functional decline up to 56 days after MI observed with IgG2a treatment (**Figure 3A**). LV volume was not different at seven days, but the dramatic increases in both diastolic and systolic volume between days seven and 56 in IgG2a-treated animals were prevented by pharmacological inhibition of CDH11 by SYN0012 (**Figure 3C-D**). This observation, combined with increased *Cdh11* transcription occurring at day seven after MI, suggests that CDH11 expressing cells play an active role in LV remodeling and dilation during the fibrotic phase of infarct healing. Albeit not statistically significant, twice as many IgG2a-treated mice died during the highest risk period for cardiac rupture (days three to seven after MI), relative to the SYN0012-treated cohort (**Figure S7**).^3^ Indeed, SYN0012 treated mice also showed increased survival relative to prior reports using this specific surgical model of MI.^14^ This difference in mortality is likely a consequence of altered tissue breakdown and inflammation-mediated softening of the myocardial wall, as measured by AFM at day seven; by day 56, we observed high stiffness variability in IgG2a treated infarcts, whereas SYN0012 treated infarcts had more consistent stiffness within the range of Sham animals (**Figure 4**).

Having identified a clear functional effect of CDH11 blockade in limiting myocardial remodeling and infarct expansion after MI, we investigated which cell types and molecular mechanisms may mediate this process. Flow cytometry-based assessment of primary non-CM cell populations in the ischemic heart revealed that the SYN0012 treatment did not affect the overall percentages of CMCs or BMDCs at three or seven days after MI (**Figure 5A-C**). However, further analysis of specific BMDC subpopulations revealed that targeting CDH11 alters the early expression of myeloid lineage macrophages, shifting the balance from inflammatory macrophages to pro-angiogenic (BMD-PACs), non-myeloid cell populations three days after MI (**Figure 5D-F**). Note that SYN0012 treatment also reduced the ratio of M1:M2-like macrophages, consistent with a pro-resolving inflammatory environment after infarct (**Figure S12**). This reduction in macrophage-like cells three days after MI may be the reason for the reduced tissue remodeling observed following SYN0012 treatment. Despite differences in the heart, the percentages of myeloid cells and BMD-PACs circulating in the peripheral blood were not significantly different three days after MI (**Figure S13**), suggesting that the SYN0012 likely has an effect on the recruitment of these cells to the heart. Alternatively, the unique mechanical environment within the heart may preferentially effect proliferation of recruited immune cells in a CDH11-dependent manner. Indeed, CDH11 deficiency has been shown to reduce MΦ recruitment, migration, proliferation, and expression of M2-MΦ markers – such as Arg1 and CD206 – in pulmonary fibrosis.^18^

We observed a reduction in *Il6* transcription with SYN0012 treatment at three days after infarct, primarily in non-CM cells (**Figure 6A**). IL-6 has been shown to have a multifaceted role in cardiovascular disease, and there is evidence that blocking IL-6 leads to worsened outcomes by reducing CM viability and angiogenesis.^19–21^ However, IL-6 has also been shown to promote both the infiltration, migration, and polarization of macrophages, and the activation of myofibroblasts.^5,22–24^ By leveraging CDH11 to selectively targeting non-CM cells in the heart, we effectively limited the negative cell activating effects of IL-6, without interfering with its function in CMs. The decrease in *Il6* transcription, and corresponding reduction in IL-6 expression in infarcted areas, was evident at three days (**Figure 6A-B**), consistent with the time course of high MΦ expression (cf. **Figure 1D-F**) and the transition from inflammatory to proliferative phase of healing. As such, there has been an increasing interest in the role of MΦs in the process of healing and cardiac remodeling after MI.^25–28^

We hypothesized that CDH11 regulates the interactions between CFs and MΦs in the heart after MI, noting that MΦ activity has been shown to alter CF function in the ischemic heart independent of CDH11.^29^ Interactions between MΦs and CFs have been reported to regulate IL-6 expression in response to TGF-β1 signaling and fibrosis.^15^ CDH11-dependent CF-MΦ interactions in the lungs have also been shown to create a self-sustaining profibrotic niche via local production of latent TGF-β1 by macrophages that can be activated by myofibroblasts.^30^ We observed increased *Il6* transcription and protein expression – a hallmark of MΦ-CF interactions – in IgG2a-treated hearts, relative to SYN0012, at three days after MI (**Figure 6A-B**). Our co-culture data confirm that interactions between MΦs and CFs promote the expression of IL-6 through a partially CDH11-dependent mechanism (**Figure 7A-B**). We also observed that transcription of *Mmp13*, a key mediator of type I collagen degradation, was increased by CF-MΦ interactions, but was not directly affected by SYN0012 treatment (**Figure 7C-D**). Therefore, we suspect that the differences in *Mmp13* transcription we observed *in vivo* – namely a faster return to Sham levels than IgG2a (cf. **Figure S14**) – may be an indirect result of the reduction in the relative abundance of MΦ subpopulations after MI. The slight but significant increase in *Tgfβ1* transcription with SYN0012 treatment in our co-culture model indicates that local expression changes may help mediate the resolution of inflammation (**Figure 7E**). TGF-β1 is a key signal to trigger the resolution of inflammation and initiation of fibrotic remodeling after MI.^31,32^

Interestingly, following CDH11 blockade we observed a reduction in transcription of both *Fgf2* and *Vegfa1*, two pro-angiogenic genes often associated with improved revascularization and better outcomes following MI.^33–36^ However, we also observed more muscularized vessels – likely arterioles – in infarcted areas of SYN0012-treated hearts at 21 days after MI; this corresponds to a time when angiogenic gene expression has run its course.^37^ We speculate that SYN0012 may either preserve native vasculature or hasten maturation of new vasculature, such that the hypoxic conditions which typically drive angiogenic signaling are reduced.^38^ Blocking CDH11 in endothelial cells could also attenuate endothelial to mesenchymal transitions, potentially contributing to preservation of the native vasculature.^8,39^ The proportion of CECs in the heart over both time points was slightly reduced with SYN0012 treatment, but increased recruitment of BMD-PACs may contribute to a restoration of vascularization (cf. **Figure 5**).

Overall, our data reveal that CDH11 engagement plays important roles in the recruitment of the BMDCs to the heart that contribute to fibrotic remodeling by activated myofibroblasts after MI. Targeting CDH11 with SYN0012 in mice does not prevent the acute inflammatory and reparative response, but rather reduces the duration of inflammation and fibrotic remodeling, resulting in a smaller, more stable infarct. We believe that this effect is mediated, in part, by limiting early recruitment of myeloid lineage BMDCs and by reducing MΦ-induced expression of IL-6 by CFs. Further, SYN0012 treatment promotes recruitment of pro-angiogenic cells (BMD-PACs) to the heart and results in the preservation of arterioles in the infarct area. Taken together, this work has identified CDH11 as a potential therapeutic target to reduce inflammatory and fibrotic remodeling following MI by selectively targeting a host of active non-CM cells responsible for the progression of cardiac fibrosis and heart failure.

## Supporting information

Compiled Supplmental Methods and Figures

## Acknowledgments

We thank the Vanderbilt Cardiovascular Physiology Core for performing echoes for this study. We thank Claire Lafferty and Joshua Bender for assisting in the collection of the AFM and histological data. We also thank Roche (Dominik Hartl & Uwe Junker, Roche Pharma Research & Early Development (pRED), Immunology, Inflammation and Infectious Diseases (I3) Discovery and Translational Area and the Cadherin-11 team, Roche Innovation Center Basel, Switzerland) for supplying the blocking antibody, SYN0012.

## Non-standard abbreviations and acronyms

αSMA: Smooth muscle α-actin
CDH11: Cadherin-11
IL-6: Interleukin-6
TGF-β1: Transforming growth factor beta 1
Fgf2: Fibroblast growth factor 2
Vegf-a: Vascular endothelial growth factor a
MMP13: Matrix metalloproteinase 13
CF: Cardiac fibroblast
Mɸ: Macrophage
CEC: Cardiac endothelial cell
CMC: Cardiac mesenchymal cell
BMDC: Bone marrow-derived cell
BMD-PAC: Bone marrow-derived pro-angiogenic cell

