## Supplementary material for "Cadherin-11 Blockade Reduces Inflammation-driven Fibrotic Remodeling and Improves Outcomes After Myocardial Infarction": Compiled Supplmental Methods and Figures

### *Detailed Methods*

#### ***Mice***

All animal procedures were approved by the Institutional Animal Care and Use Committee at Vanderbilt University. Myocardial infarction (MI) was induced by permanent ligation of the left anterior descending (LAD) coronary artery, as previously described (Gao et al. 2010). Briefly, mice were anesthetized by 2% isoflurane inhalation, without ventilation, and a small skin incision was made over the left chest. After dissection and retraction of the pectoral major and minor muscles, the 4th intercostal space was exposed and a small hole was made to open the pleural membrane and pericardium. The heart was then gently popped out through the hole and the LAD was ligated ~3 mm from its origin using a 6-0 silk suture. Successful ligation was confirmed by pale coloration of the anterior wall of the left ventricle. Following LAD ligation, the heart was placed back into the intra-thoracic space, air was evacuated from the thoracic cavity, and the muscle and skin was sutured closed. MI was performed in 12 to 16 week old male *Cadherin-11* transgenic mice ($Cdh11^{+/+}$, $Cdh11^{+/-}$, $Cdh11^{-/-}$) or WT mice (C57BL/6J; Jackson Laboratories). All mice were given pre- and post-operative analgesic of 5 mg/kg ketoprofen every 24 hours for 72 hours.

For antibody treatments, mice were administered either 10 mg/kg of a cadherin-11 (CDH11) functional blocking antibody (SYN0012; with permission from Roche) or an isotype control antibody (IgG2a) resuspended in sterile saline. Antibodies were delivered by intraperitoneal (IP) injection every four days, beginning one day after surgery, with the last treatment given on day 17 after infarct.

#### ***Isolation of cardiomyocyte and non-cardiomyocyte cells***

Live cardiac cells were isolated from mouse hearts as previously described (O'Connell, Rodrigo, and Simpson 2007). Briefly, slow perfusion of the heart with a collagenase solution to digest the ECM and isolate intact cardiomyocytes (CMs) and non-CMs from hearts. Cells were separated into CM and non-CM fractions by centrifugation for 10 min at 90 g and lysed in TRIZOL for analysis of *Cdh11* transcription.

#### ***Identification of non-CM cell types by flow cytometry***

Composition of cell types in the heart and the peripheral blood were measured by flow cytometry. Hearts were isolated from animals at three or seven days after Sham or MI surgeries and immediately placed in a solution of ice cold FACS buffer (5% FBS in PBS). The atria were removed and the ventricular tissue was minced and stored on ice. Minced samples were placed in 2 mL of digestion solution, comprised of 1.4 mg/mL of type II collagenase in HBSS, and incubated at 37°C for seven minutes. The digested sample was then filtered through a 100 μm cell strainer and resultant cells were washed in 50 mL of ice cold FACS buffer, centrifuged at 1500 rpm for 5 minutes, resuspended in FACS buffer and filtered through a 70 μm cell strainer into room temperature red blood cell lysis buffer (BioLegend). Similarly, peripheral blood was collected with a K-EDTA syringe by cardiac puncture and quickly diluted into room temperature red blood cell lysis buffer. After 5 minutes in lysis buffer, cells were washed in FACS buffer, centrifuged at 1500 rpm for 5 minutes, and counted to measure the total number of cells. Cells were then taken from each sample, suspended in DAPI (1:100,000; Thermo-Fisher Scientific) to identify dead cells, and stained with conjugated antibodies for Ter-119 (1:100; violetFluor™ 450 clone TER-119; Tonbo Biosciences), CD45.2 (1:100; PerCP-Cy5.5 clone 104; Tonbo Biosciences), CD31 (1:100; PE-Cy7 clone MEC13.3; BioLegend), CD11b (1:100; PE clone M1/70; eBioscience), CD206 (1:100; APC clone MR6F3; eBioscience), CD86 (1:100; FITC clone GL1; eBioscience), Gr-1 (1:100; FITC clone RB6-8C5; eBioscience). Staining for CDH11 was performed in a two-step process with an unconjugated primary antibody (1:100; clone 23C6; from M. Brenner) followed by a PE secondary antibody (1:100; PE clone; RMG1-1; BioLegend). Note that antibodies specific for CD45.1 (1:100; PE clone A20; BD Biosciences) and CD45.2 (1:100; FITC clone 104; BD Biosciences) were used for assessment of bone marrow engraftment efficiency (**Figure S6**).

Gating based on size and shape (FSC and SSC), in addition to negative staining for DAPI and Ter-119, was used to identify viable single cells (**Figure S1**). Further gating based on fluorescence minus one (FMO) stained controls was used to identify the presence of distinct cell populations (**Figure S2**). In particular, we identified bone marrow derived cells (BMDC: $CD45^{+}$), cardiac endothelial cells (CEC: $CD45^{-}CD11b^{-}CD31^{+}$), and cardiac mesenchymal cells – primarily myofibroblasts – (CMC: $CD45^{-}CD11b^{-}CD31^{-}$). Within the BMDC population, we gated for bone marrow-derived proangiogenic cells (BMD-PAC: $CD45^{+}CD11b^{-}CD31^{+}$) and myeloid lineage cells ($CD45^{+}CD11b^{+}$). Within the myeloid cell population, we assessed macrophage polarization by gating for pro-inflammatory macrophages (M1-like MΦ: $CD45^{+}CD11b^{+}CD86/Gr\text{-}1^{+}CD206^{-}$) and pro-healing macrophages (M2-like MΦ: $CD45^{+}CD11b^{+}CD86/Gr\text{-}1^{+}CD206^{+}$) (**Figure S1**). Using the FMO control for CDH11 (**Figure S2**), this gating strategy was applied to 1) all live, single cells and 2) all live, CDH11+ cells (**Figure S4C-D**). In this manner, we were able to compute the fraction of CDH11+ cells in each cell population of interest within the heart (**Figure S3A** and **Figure 1**) and peripheral blood (**Figure S3B** and **FigureS5**). For studies involving SYN0012 treatment (**Figure 5** and **Figure S13**), the same gating strategy was used to identify cell populations of interest, although staining for CDH11 expression was not performed.

#### *Quantification of cardiac function and geometry by echocardiography*

Ejection fraction (EF), left ventricular (LV) mass, and LV volume were measured from short axis cardiac M-Mode images captured on the Vevo 2100 small animal ultrasound system (VisualSonics). A minimum of 6 independent measures of LV diameter and wall thickness were used to calculate metrics of cardiac function and geometry for each mouse at each time point. Note that EF, LV mass and LV volume were calculated from the LV inner diameter at diastole and systole.

Measurements were made just prior to MI, as well as at days seven, 21, and 56 days after surgery. Mice with an EF reduced by less than 5% or greater than 60% were excluded from subsequent analyses. This ensured that all mice included in the study had consistently-sized (intermediate to large) infarcts which had not progressed to complete heart failure. Mice were euthanized by $CO_2$ inhalation in accordance with university guidelines at three, seven, 21 and 56 days after infarct for further processing.

#### *Bone marrow transplantation*

6 week old male WT mice expressing the CD45.1 allele (B6.SJL-*Ptprc^a^Pepc^b^*/BoyJ; Jackson Laboratories) were lethally irradiated with a 10 Gy split dose from a $Cs^{137}$ source (5 Gy in the morning followed by 5 Gy 4-6 hours later). Within 24 hours, age- and gender-matched *Cdh11*$^{+/+}$, *Cdh11*$^{+/-}$, and *Cdh11*$^{-/-}$ donors were euthanized and bone marrow was isolated from whole femurs. Donor bone marrow was delivered to irradiated recipients by retro-orbital injection ($1\times10^6$ cells in 100μL), and recipient mice were maintained on acidified water with antibiotics (neomycin and polymyxin B) for up to 2 weeks. Transplant efficiency was confirmed by flow cytometric analysis of isolated bone marrow showing simultaneous expression of the donor CD45.2 allele and absence of the original CD45.1 allele (**Figure S6**). To allow sufficient time for bone marrow reconstitution, mice received MI by permanent LAD ligation six weeks after transplantation.

#### *Cryosectioning and quantitative histological analysis*

Following euthanasia, hearts were dissected into PBS, weighed (**Figure S10A-B**), and then submerged briefly in a KCl solution to relax the CMs. Relaxed hearts were then bisected along the transverse plane (orthogonal to the long axis of the heart), embedded in OCT media, and frozen. Frozen blocks were subsequently cryosectioned into 10 μm sections, mounted onto glass slides, and stored at -20°C. A selection of the slides were stained using Masson's trichrome (Sigma), according to the manufacturer's instructions, in order to identify regions of healthy myocardium (red/pink), collagen/ECM deposition (blue), and cell nuclei (black) (**Figure S9**). Prior to staining, slides were brought to room temperature, OCT media was dissolved in PBS, and sections were fixed in Bouin's solution.

To quantify infarct morphology (length and thickness) from Masson's trichrome stained sections, we developed a semi-automated image processing pipeline (**Figure S9**). Briefly, non-uniformity in background illumination was corrected by bottom hat filtering prior to boundary detection. Using the detected boundaries, local ventricular wall thickness was computed using an Eulerian solution to a pair of linear partial differential equations over the histological domain (Yezzi and Prince 2003). By solving the Laplace equation for a given harmonic function $f$ between the inner (ventricular lumen) and outer (myocardial wall) boundaries, the local thickness field was obtained. Correspondence trajectories along $f$ between the inner and outer boundaries were used to subdivide histological sections into 40 circumferential partitions. (Rocha, Yezzi, and Prince 2007). Following partitioning, colorimetric segmentation in an HSL color space (Bersi et al. 2017) was performed in order to identify the area fractions of myocardium (red; H = 250° - 25°, S = 0.1 - 1.0, L = 0.1 - 0.93) and collagen (blue; H = 150° - 250°, S = 0.1 - 1.0, L = 0.1 - 0.93) within each partition. Based on the identified area fractions, locations of infarct borders were defined based on the inversion of area fractions (i.e., clockwise from red > blue to blue > red and vice-versa). Using this approach, the location and thickness profile of the infarcted regions were automatically identified. In cases of poor automatic detection, infarct borders were defined manually. Finally, following identification of infarct borders, the ratio of infarct length to total circumferential length (in degrees) was used to estimate the percent circumference of the infarct region. Using the thickness profile between infarct borders, the average and standard deviation (i.e., thickness variation) were used to characterize infarct morphology (**Figure S10D-E**). Quantification was performed on 3 sections per heart separated by at least 300 μm.

#### *Immunohistochemistry*

Frozen slides were brought to room temperature, OCT media was dissolved in PBS, and tissue sections were fixed in 4% paraformaldehyde with 0.3% Triton-X for 10 min followed by blocking in 1% BSA in PBS for 1 hour. Tissue sections were then stained for either αSMA (Sigma/BD Biosciences), CD31 (Biolegend), and IL-6 (Abcam). Sections stained with non-conjugated antibodies (IL-6) were incubated at a 1:100 dilution in 1% BSA overnight at 4°C. Sections were then rinsed with PBS and incubated with fluorescently tagged secondary antibodies for 1 hour at a 1:300 dilution in 1% BSA. Sections stained with directly conjugated antibodies were incubated at a 1:100 dilution in 1% BSA for 1 hour at room temperature. Stained slides were mounted in ProLong Gold with DAPI to visualize cell nuclei and were imaged using an Olympus BX53 microscope equipped with a high resolution Qimaging Retiga 3000 camera.

#### *Atomic force microscopy*

Frozen slides were acclimated to room temperature, OCT media was dissolved in PBS, and tissue sections were blocked in 10% FBS for 20 minutes. Tissue sections were then stained for αSMA (Sigma) and a Hoechst nuclear stain (Invitrogen) for 20 minutes to allow for visualization of the infarct while scanning with the atomic force microscope (AFM). The Biocatalyst AFM developed by Bruker was used to measure tissue topography and stiffness within SYN0012 and IgG2a treated infarcts. The AFM probe was equipped with a blunted pyramidal tip specifically developed for soft biological samples (MLCT-Bio) and the peak force quantitative nanomechanical mapping scanning mode (PeakForce QNM) was used in order to provide robust measurements of topography and elastic modulus. Prior to scanning tissue samples, the system was calibrated on 40 kPa polyacrylamide gel standards and the spring constant and deflection sensitivity of the AFM probe was calculated. All measurements were made in PBS and were acquired from at least 5 separate 10x10 $\mu m^2$ areas from a minimum of 2 different sections per mouse (cf. **Figure S8**).

#### *Cell isolation and culture*

To complement and inform our *in vivo* studies, cardiac fibroblasts (CFs) (Golden et al. 2012) and intraperitoneal macrophages (MΦs) were isolated from mice. CFs were isolated from *Cdh11*$^{+/+}$ and *Cdh11*$^{-/-}$ mice bred onto the *Immorto* mouse line, such that cells from littermate controls could be maintained in culture for longer. Briefly, hearts from 8 week old mice were isolated, minced, and digested in 2% collagenase solution supplemented with trypsin for the last 10 minutes of a 40 minute digest. CFs were then rinsed with PBS, transferred to gelatin coated plates, and cultured in DMEM supplemented with 10% FBS, 1% penicillin/strep, and interferon gamma at 33°C in order to maintain the immortalized phenotype. Prior to experimental use, cells were replated in DMEM supplemented with 10% FBS and 1% penicillin/strep and grown at 37°C for 48 hours to deactivate the immortalized gene.

MΦ exfiltration was stimulated by intraperitoneal (IP) injection of 1mL of 4% thioglycollate media into C57 black 6 mice. After 72 hours, mice were sacrificed and the intraperitoneal cavity flushed with 10 mLs of cold RPMI media to collect the cells. After washing in cold PBS, cells were plated in RPMI media supplemented with 10% FBS on tissue culture plastic and allowed to adhere for 1 hour. Following previous reports, non-adherent cells were then rinsed away and all remaining cells were taken to be as MΦs (Zhang, Goncalves, and Mosser 2008).

#### *Gel contraction assay*

CFs (WT and *Cdh11*$^{-/-}$) were diluted in a 1.28 mg/mL collagen solution derived from PureCol (Advanced Biomatrix) to a final concentration of 250 thousand cells/mL and were poured into a Teflon ring in a suspension well. After polymerizing for 1 hour, DMEM supplemented with 10% FBS, 1% penicillin/strep was added to flood the well and release the collagen gel from both the bottom of the well and the Teflon ring. Gels were imaged immediately after release and at multiple times over the next 48 hours. At each time point, gel areas were measured in ImageJ and normalized to the original gel area. For comparison of the impact of IgG2a and SYN0012 treatment, antibody was added to the cell/gel mixture at a final concentration of 20 μg/ml prior to pouring; media added to the well also contained 10 μg/ml of antibody (**Figure S11**)

#### *qPCR*

For assessment of *in vivo* transcription of specific profibrotic and inflammatory genes of interest, hearts were isolated under RNAse-free conditions and immediately flash frozen. For isolation of mRNA, samples were subsequently thawed and lysed in TRIZOL with chloroform induced phase separation according to manufacturer's instructions. cDNA was synthesized using the Superscript IV kit (Invitrogen) using 500 ng of mRNA. Real time qPCR was used to amplify targets from the cDNA using a SYBR green master mix (BIO-RAD) and specific primer sets (**Table S1**). The BIO-RAD CFX96 C1000 system was used to quantify gene transcription in each sample, relative to *Gapdh*. For all of the *in vivo* transcription levels, post-MI samples were normalized to the average of all three and seven day Sham values.

**Table S1** qPCR primers

| Target | Forward | Reverse |
|---|---|---|
| ***Gapdh*** | ATGACAATGAATACGGCTACAG | TCTCTTGCTCAGTGTCCTTG |
| ***Ctnt*** | AGGAGCTGATTTCCCTCAAAG | TTTCCTTCTCCCGCTCATTG |
| ***Cdh11, exon12*** | TCACTATCAAAGTCTGTGGCTG | CAAACAGCACAACGATGACC |
| ***Il6*** | CAAAGCCAGAGTCCTTCAGAG | GTCCTTAGCCACTCCTTCTG |
| ***F4/80*** | ACCACAATACCTACATGCACC | AAGCAGGCGAGGAAAAGATAG |
| ***Il1β*** | TCCTGTGTAATGAAAGACGGC | ACTCCACTTTGCTCTTGACTTC |
| ***Tnfα*** | AGACCCTCACACTCAGATCA | TGTCTTTGAGATCCATGCCG |
| ***Mmp3*** | CAGGAAGATAGCTGAGGACTTTC | GGTCAAATTCCAACTGCGAAG |
| ***Mmp13*** | GATTATCCCCGCCTCATAGAAG | TCTCACAATGCGATTACTCCAG |
| ***Tgfβ1*** | CCTGGGTTGGAAGTGGATC | TTGGTTGTAGAGGGCAAGG |
| ***αsma*** | GAGAAGCCCAGCCAGTCG | CTCTTGCTCTGGGCTTCA |
| ***Col1a1*** | CACCCTCAAGAGCCTGAGTC | GTTCGGGCTGATGTACCAGT |
| ***Fgf2*** | GGAGTTGTGTCTATCAAGGGAG | TGCCCAGTTCGTTTCAGTG |
| ***Vegfa1*** | AAAGCCAGCACATAGGAGAG | CGAGTCTGTGTTTTTGCAGG |
| ***Mrc1*** | ATGGATGTTGATGGCTACTGG | TTCTGACTCTGGACACTTGC |
| ***Cd14*** | CCTTTCTCGGAGCCTATCTG | CAACTTTCCTCGTCTAGCTCG |
| ***Arg1*** | AAGAATGGAAGAGTCAGTGTGG | GGGAGTGTTGATGTCAGTGTG |

***Quantification of IL-6 production by indirect ELISA***

Fifty thousand CFs were plated in a 12 well plate and allowed to adhere for 20 min prior to exposure to media containing between 0 and fifty thousand MΦs (final volume of 1.3 mL per well). We tested the interaction of cells (including MΦs alone) without antibody treatment, and then specifically compared CFs and a range of co-culture with IgG2a or SYN0012. For antibody treatments, samples were incubated with antibody (10 μg/ml) for 15 minutes before plating. After 48 hours in culture, conditioned media was removed from each well and IL-6 secretion was measured with a DuoSet mouse IL-6 ELISA (R&D Systems). After boiling, 100 μl of each sample was added in duplicate and compared against a provided standard (**Figure 7A-B**). Cells from these co-cultures were also lysed in TRIZOL for isolation of mRNA, cDNA synthesis, and analysis of transcription of specific gene targets by qPCR, as described above (**Figure 7C-E** and **Figure S15**).

***Statistical analysis***

For all experiments measuring outputs across a range of time points and treatments, a two-way ANOVA was run to determine significant effects of time, treatment, and interactions between the two. The post-hoc Holms-Sidak method and individual student t-tests with an overall significance level of 0.05 were used to account for multiple comparisons within cell types and treatment groups. Non-parametric tests (ANOVA on ranks or rank sum tests) were used if samples either failed the Shapiro-Wilks normality test or had unequal variance. F-tests were run on median stiffness values from AFM in order to compare variances of tissue stiffness. For all comparisons, a value of $p < 0.05$ was considered significant and differences have been indicated in figures throughout the manuscript and supplement, where appropriate.

***References***

Bersi, M R, R Khosravi, A J Wujciak, D G Harrison, and J D Humphrey. 2017. “Differential Cell-Matrix Mechanoadaptations and Inflammation Drive Regional Propensities to Aortic Fibrosis, Aneurysm or Dissection in Hypertension.” *Journal of the Royal Society, Interface* 14 (136):1–31. https://doi.org/10.1098/rsif.2017.0327.

Gao, Erhe, Yong Hong Lei, Xiying Shang, Z. Maggie Huang, Lin Zuo, Matthieu Boucher, Qian Fan, J. Kurt Chuprun, Xin L. Ma, and Walter J. Koch. 2010. “A Novel and Efficient Model of Coronary Artery Ligation and Myocardial Infarction in the Mouse.” *Circulation Research* 107 (12). Lippincott Williams & WilkinsHagerstown, MD:1445–53. https://doi.org/10.1161/CIRCRESAHA.110.223925.

Golden, H B, D Gollapudi, F Gerilechaogetu, J Li, R J Cristales, X Peng, and D E Dostal. 2012. “Isolation of Cardiac Myocytes and Fibroblasts from Neonatal Rat Pups.” *Methods Mol Biol* 843:205–14. https://doi.org/10.1007/978-1-61779-523-7_20.

O’Connell, Timothy D., Manoj C. Rodrigo, and Paul C. Simpson. 2007. “Isolation and Culture of Adult Mouse Cardiac Myocytes.” In *Cardiovascular Proteomics*, 271–96. New Jersey: Humana Press. https://doi.org/10.1385/1-59745-214-9:271.

Rocha, Kelvin R., Anthony J. Yezzi, and Jerry L. Prince. 2007. “A Hybrid Eulerian-Lagrangian Approach for Thickness, Correspondence, and Gridding of Annular Tissues.” *IEEE Transactions on Image Processing* 16 (3):636–48. https://doi.org/10.1109/TIP.2007.891072.

Yezzi, Anthony J, and Jerry L Prince. 2003. “An Eulerian PDE Approach for Computing Tissue Thickness.” *IEEE Transactions on Medical Imaging* 22 (10):1332–39. https://doi.org/10.1109/TMI.2003.817775.

Zhang, X, R Goncalves, and D M Mosser. 2008. “The Isolation and Characterization of Murine Macrophages.” *Curr Protoc Immunol* Chapter 14:Unit 14 1. https://doi.org/10.1002/0471142735.im1401s83.

*Supplemental Figures*

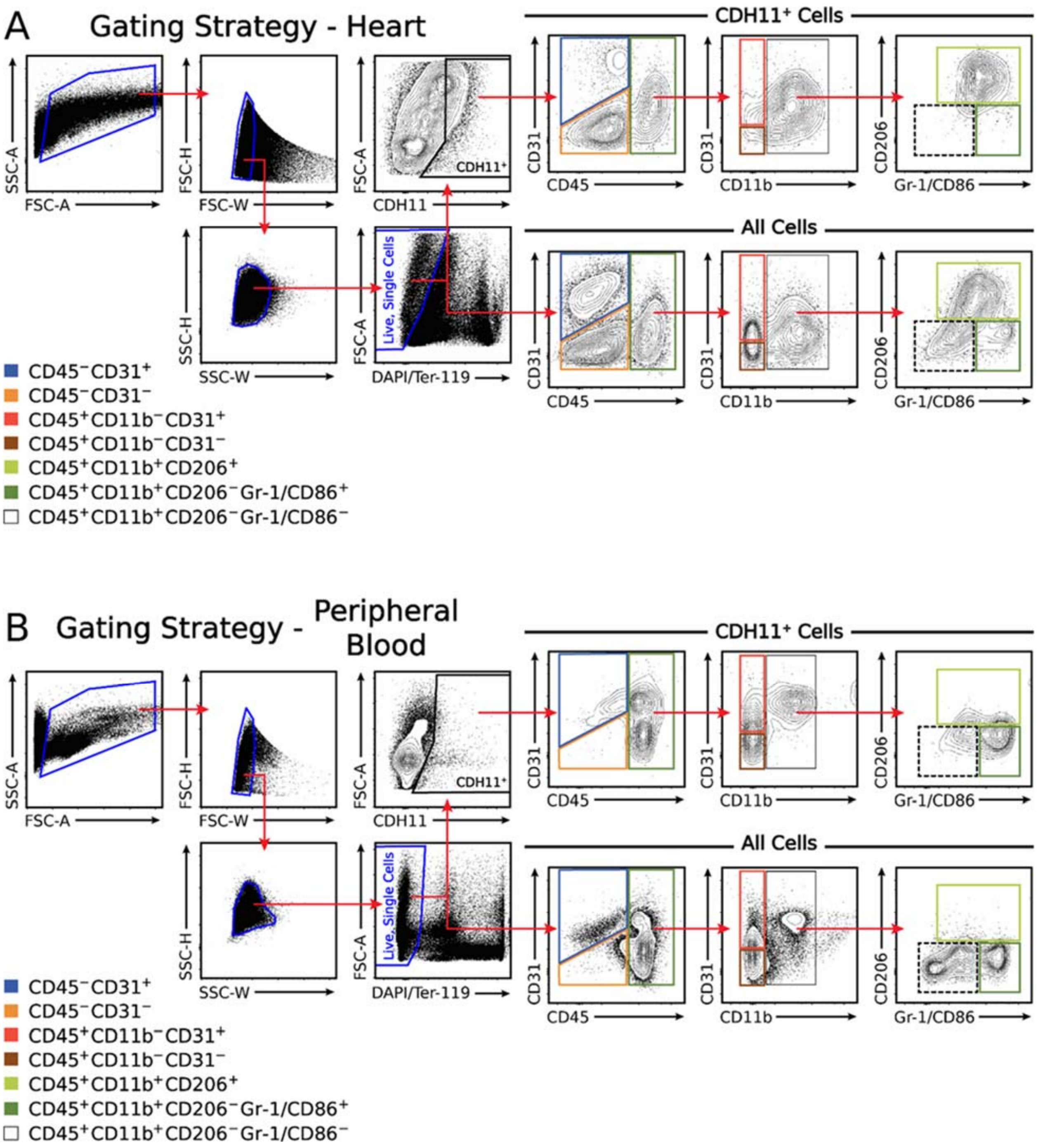


**Figure S1. Flow cytometry gating strategy for primary cell populations of interest.** Live, single cells were identified from all measured events in the heart (**A**) and peripheral blood (**B**). Cells were separated into cardiac endothelial cells (blue), cardiac mesenchymal cells (orange), and bone-marrow derived cells. BMDCs were further gated into bone marrow-derived proangiogenic cells (red), non-myeloid lineage BMDCs (brown), and myeloid lineage cells which were further gated into M1-like macrophages (dark green), M2-like macrophages (light green), and a remaining myeloid subset (white/dashed). Gating was applied to all events (bottom rows) and CDH11$^+$ cells (top rows) and is shown for representative samples at 7 days post-MI.

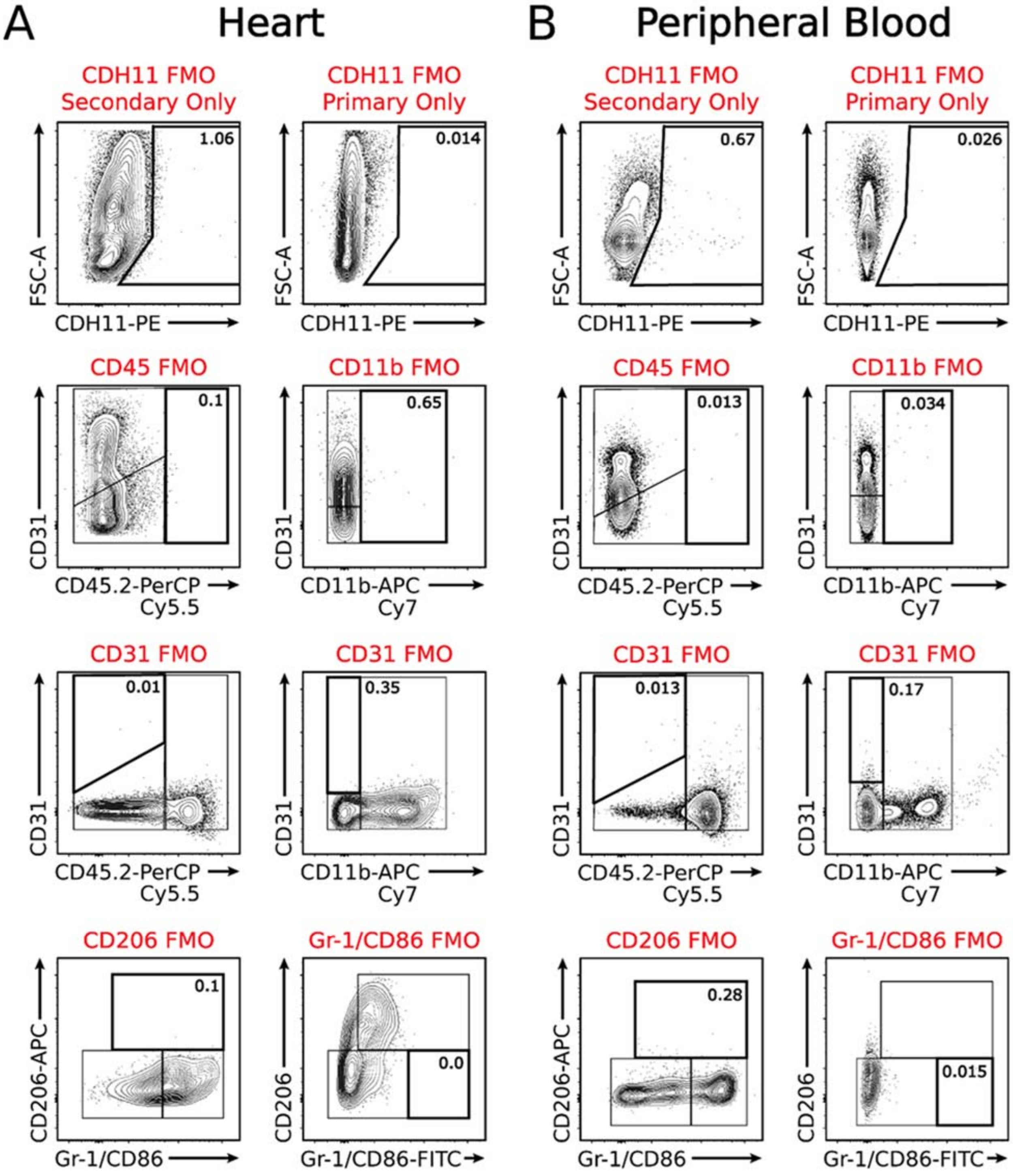


**Figure S2. Fluorescence minus one (FMO) controls for flow cytometry analysis.** FMO controls for each antibody used in the flow cytometric analyses indicates the placement of gates defining each of the primary cell populations in the heart (**A**) and peripheral blood (**B**). Inset numbers show the percent of parent population events that fall within each gate. Note that gates were placed such less than ~1% of events occurred within the positive gate.

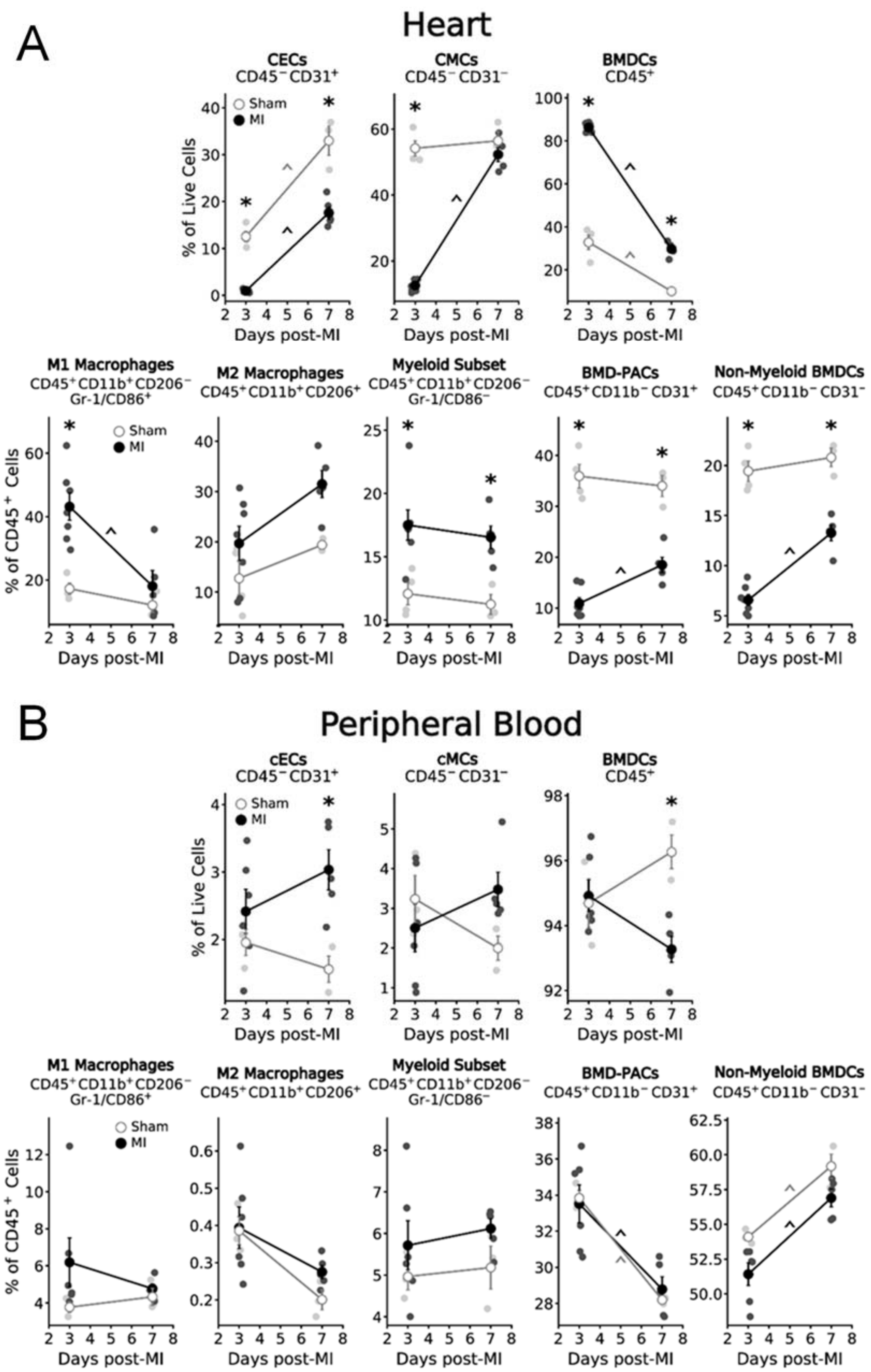


**Figure S3. Resident and bone marrow derived cell expression post-MI.** Flow cytometric analysis reveals the percentage each of the identified cell populations, relative to total live single cell events, in the heart (**A**) and peripheral blood (**B**) after MI. * $p < 0.05$ between Sham and MI at the same time, ^ $p < 0.05$ over time; n = 3-7 per group.

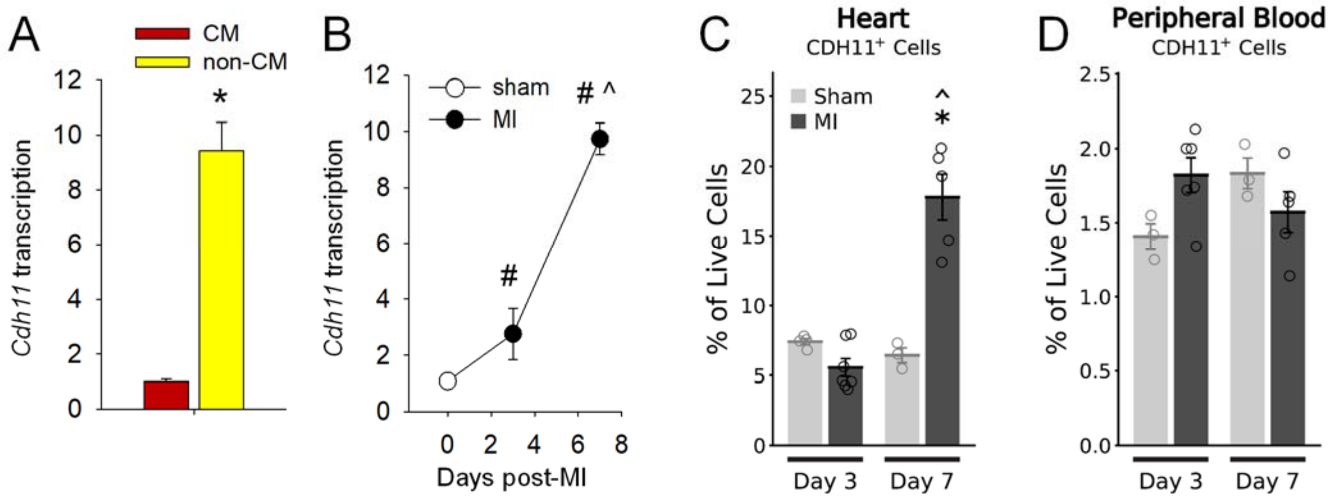


**Figure S4. CDH11 expression post-MI.** Transcription of *Cdh11* occurs primarily in non-CM cells (**A**) and increases up to maximal expression (~10-fold higher than Sham) at day seven after MI (**B**). Flow cytometric analysis reveals the percentage of $CDH11^+$ cells, relative to total live single cell events, in the heart (**C**) and peripheral blood (**D**) after MI. For **A-B**: * $p < 0.05$ between cell types, # $p < 0.05$ relative to Sham, ^ $p < 0.05$ relative to previous time point; For **C-D**: * $p < 0.05$ between Sham and MI at the same time, ^ $p < 0.05$ over time; n = 3-7 per group.

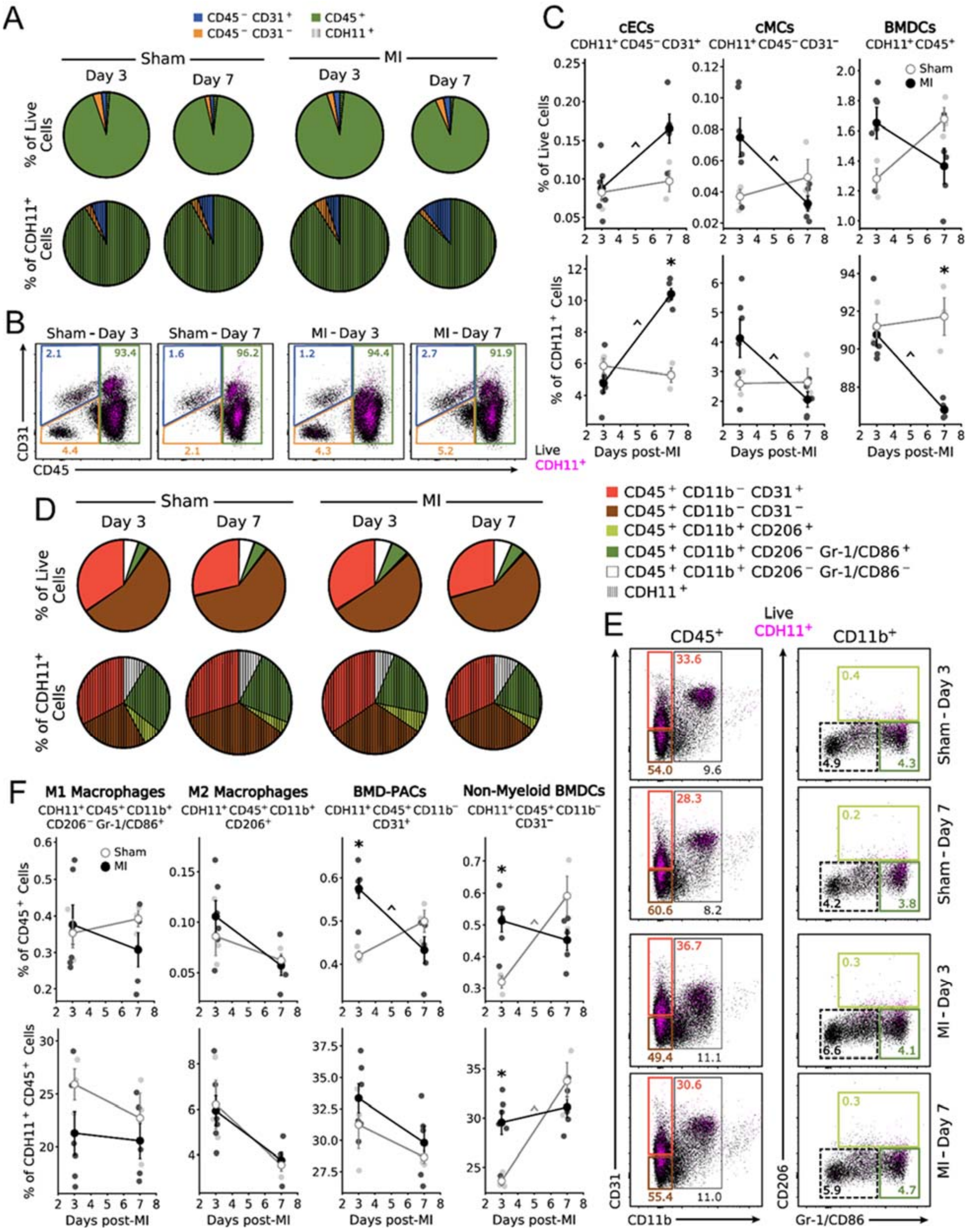


**Figure S5. CDH11 expression in the blood is largely unaffected post-MI.** Flow cytometric analysis reveals that circulating endothelial (cEC), mesenchymal (cMC), and bone marrow derived (BMDC) cell populations have minimal baseline CDH11 expression and are largely unaffected post-MI (hatched wedges). Pie charts (**A**) are scaled by either total number of live single cells (top row) or total number of CDH11 expressing cells (bottom row), relative to Sham at day three. Representative dot plots (**B**) show CDH11 expression (magenta) within each cell population (colored gates). CDH11 positive cells (**C**) within each population are shown as either a percentage of live cells or a percentage of all CDH11 positive cells. While CDH11 expression was low (<2% of total cells), the percent of cECs expressing CDH11 is increased and the percent of BMDCs expressing CDH11 is decreased, relative to Sham, at day seven post-MI. BMDC subpopulations (**D**) revealed little CDH11 expression with little difference between Sham and MI. Representative dot plots (**E**) show CDH11 expression (magenta) within each subpopulation (colored gates). Though CDH11 expression was low in each BMDC subpopulation (<1% of $CD45^+$ cells), there was an increase in $CDH11^+$ BMD-PACs and $CDH11^+$ non-myeloid BMDCs at three days post-MI, with non-myeloid BMDCs comprising significantly more of all CDH11+ BMDCs than Sham at day three. * $p < 0.05$ between Sham and MI at the same time, ^ $p < 0.05$ over time; n = 3-7 per group.

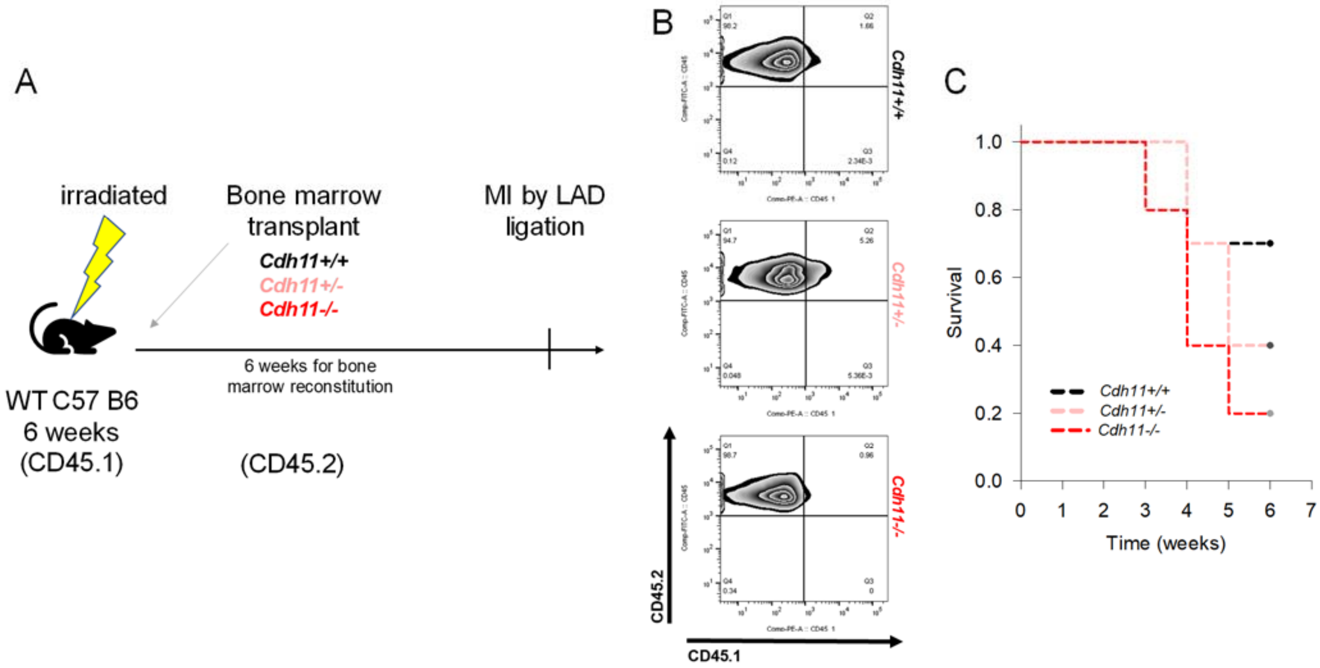


**Figure S6: Experimental protocol for bone marrow transplantation.** Schematic of experimental protocol for bone marrow transplantation from *Cdh11* transgenic donors (CD45.2) into irradiated C57BL6/J recipients (CD45.1; B6.SJL-*Ptprc*$^{a}$*Pepc*$^{b}$/BoyJ) (**A**). Flow cytometry of CD45.1 and CD45.2 expression in the bone marrow after 6 weeks of reconstitution used to confirm successful bone marrow transplantation (**B**). Despite successful transplantation, fewer mice with *Cdh11* transgenic bone marrow (e.g., *Cdh11*$^{+/-}$ and *Cdh11*$^{-/-}$) survived to the point of MI (**C**).

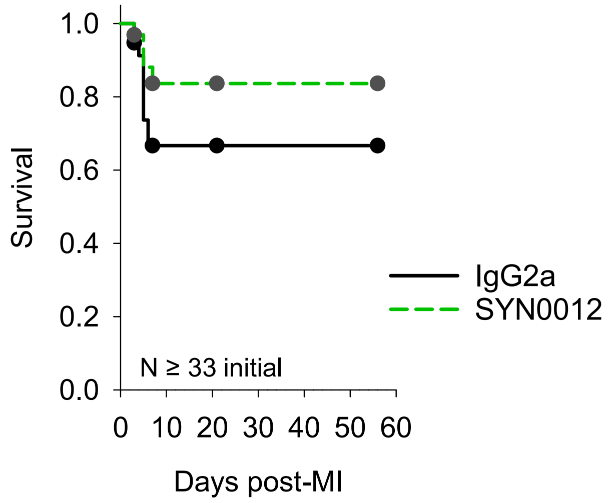


**Figure S7: CDH11 blockade improves survival after MI.** Following MI, fewer animals receiving SYN0012 treatment died as compared to those receiving IgG2a. Though survival was improved, the findings were not statistically significant with a value of $p = 0.16$.

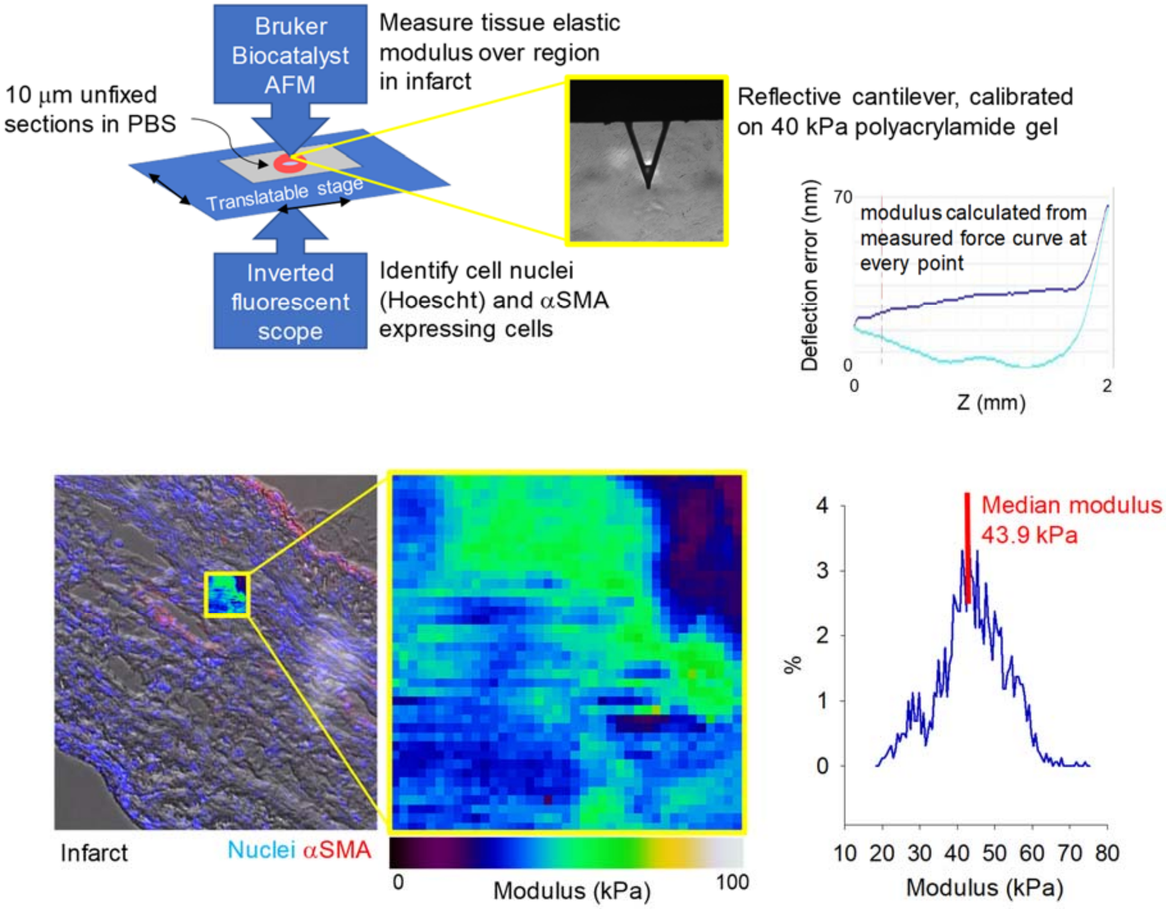


**Figure S8: Atomic force microscopy technique.** Schematic of AFM/microscope arrangement, with callout image of pyramidal probe cantilever in top panel (yellow). Typical indentation curve from PeakForce QNM mode shown at top right. Representative scan of 10x10$\mu m^2$ infarct tissue with nuclei in blue and αSMA in red. Inset shows stiffness (modulus) colormap and histogram of shows the distribution and median of measured moduli in lower panel.

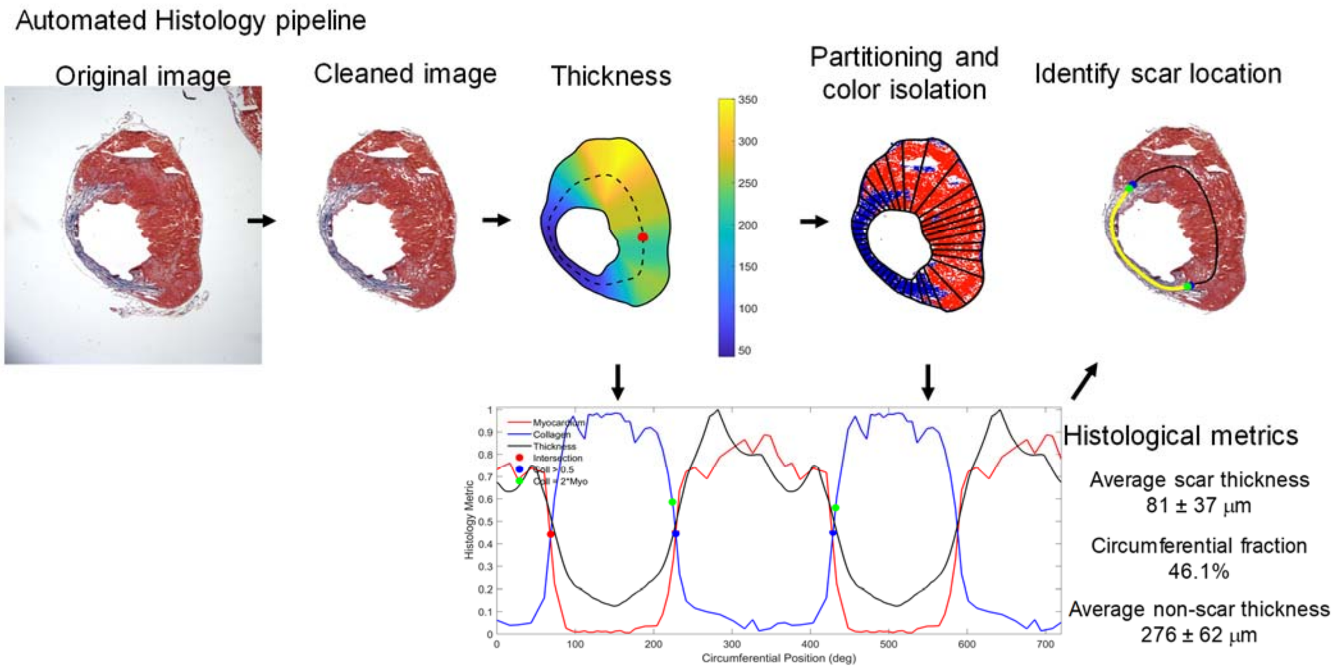


**Figure S9: Semi-automatic histological image analysis pipeline.** Masson's trichrome stained short-axis sections of infarcted hearts were analyzed in order to quantify infarct morphology (i.e., circumferential percentage and thickness). Following background removal, thickness between inner and outer boundaries is computed, 40 circumferential partitions are defined, and colorimetric segmentation of red and blue pixels is performed (shown as pseudo-colored image). Infarct borders are defined based on intersections between red/blue area fractions (bottom panel); infarct and non-infarct morphology is computed following identification of infarct borders.

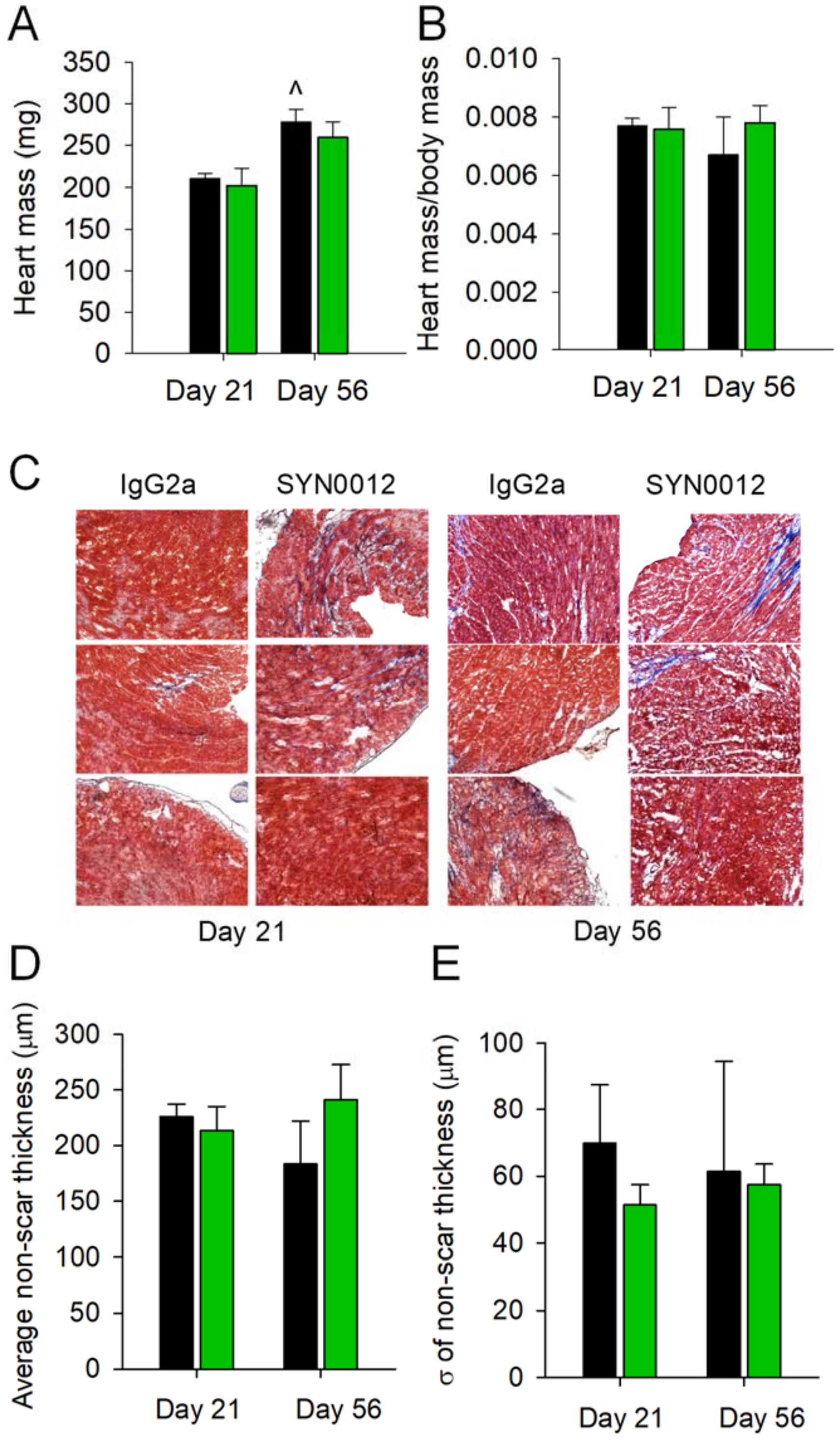


**Figure S10. Measurements of cardiac hypertrophy and fibrosis.** Comparison between IgG2a and SYN0012 treated heart mass (**A**) heart to body mass ratio (**B**), and remote interstitial fibrosis (**C**) at days 21 and 56. There was also no difference in the measured average and standard deviation ($\sigma$) of the non-scar region thickness (**D-E**). N $\geq$ 3; ^ $p < 0.05$ between timepoints

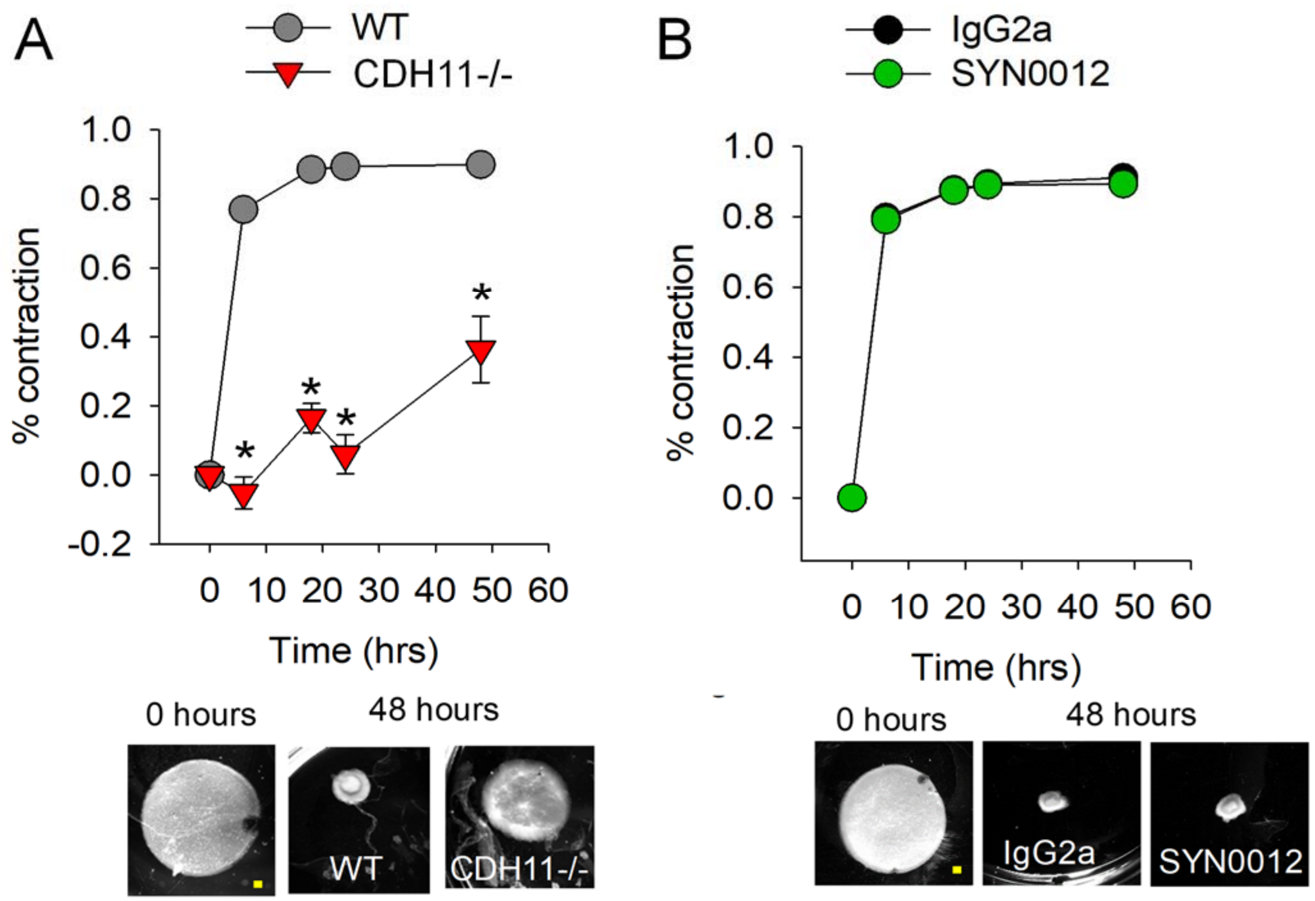


**Figure S11: CDH11 affects cardiac fibroblast (CF) contraction.** CFs isolated from *Cdh11*$^{-/-}$ mice exhibit a reduction in contractile ability (i.e., reduced gel contraction) as compared to *Cdh11*$^{+/+}$ CFs (**A**). Treatment of WT (C57BL6/J) CFs with a CDH11 blocking antibody (SYN0012) does not affect CF contractility and gel contraction (**B**). Yellow scale bar represents 1 mm. n = 3 per group.

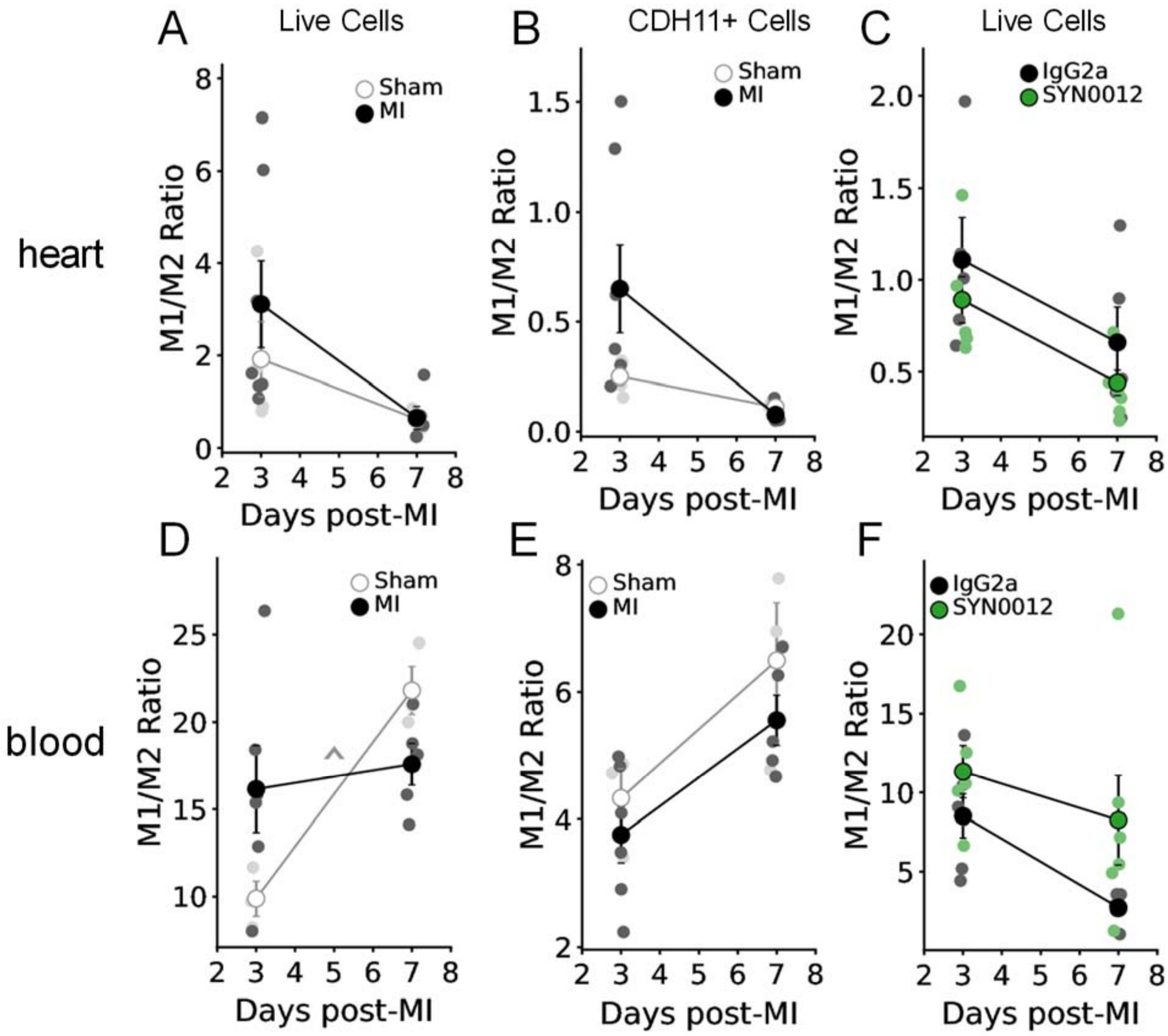


**Figure S12. Ratios of M1:M2-like macrophages post-MI.** Flow cytometric analysis reveals that the ratio of M1:M2-like macrophages in the heart was not significantly different between day three and day seven when compared between all live cells (**A**), CDH11$^{+}$ cells (**B**), or live cells treated with IgG2a or SYN0012 (**C**). Though not statistically different, SYN0012 treatment led to a 19.8% and 33.3% reduction in the M1:M2 ratio at day three and seven after infarct, respectively. The M1:M2 ratio of live cells in the peripheral blood significantly increased in Sham animals between day three and seven (**D**), but was not significantly affected in the CDH11$^{+}$ populations (**E**) or by SYN0012 treatment (**F**).

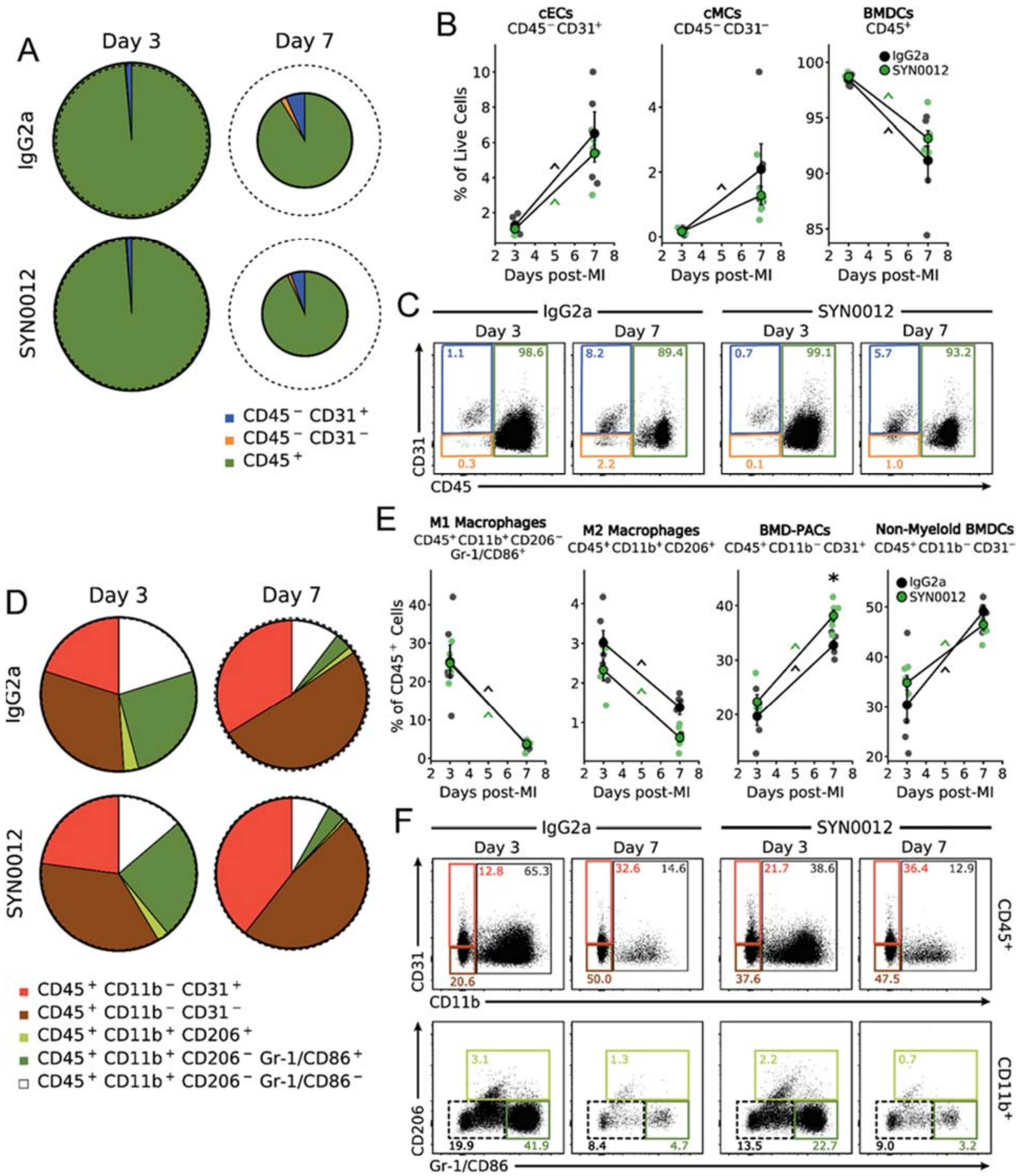


**Figure S13. CDH11 blockade has little effect on circulating cell populations post-MI.** Flow cytometric analysis reveals that CDH11 blockade by SYN0012 does not alter the percentages of circulating endothelial (cEC), mesenchymal (cMC), and bone marrow derived (BMDC) cells in the blood post-MI, relative to IgG2a (**A**). Dotted circles denote the number of cells in Sham blood samples at three and seven days after infarct. Differences are seen over time and there is a trend toward decreased cMCs and increased BMDCs after SYN0012 treatment (**B**). Representative dot plots (**C**) show changes in expression of each cell population (colored gates). Separation of BMDC subpopulations (**D**) revealed that SYN0012 significantly increases circulating BMD-PACs at day seven in addition to changes over time (**E**). Representative dot plots (**F**) show changes in expression of each BMDC subpopulation (colored gates). * $p < 0.05$ between Sham and MI at the same time, ^ $p < 0.05$ over time; n = 3-7 per group.

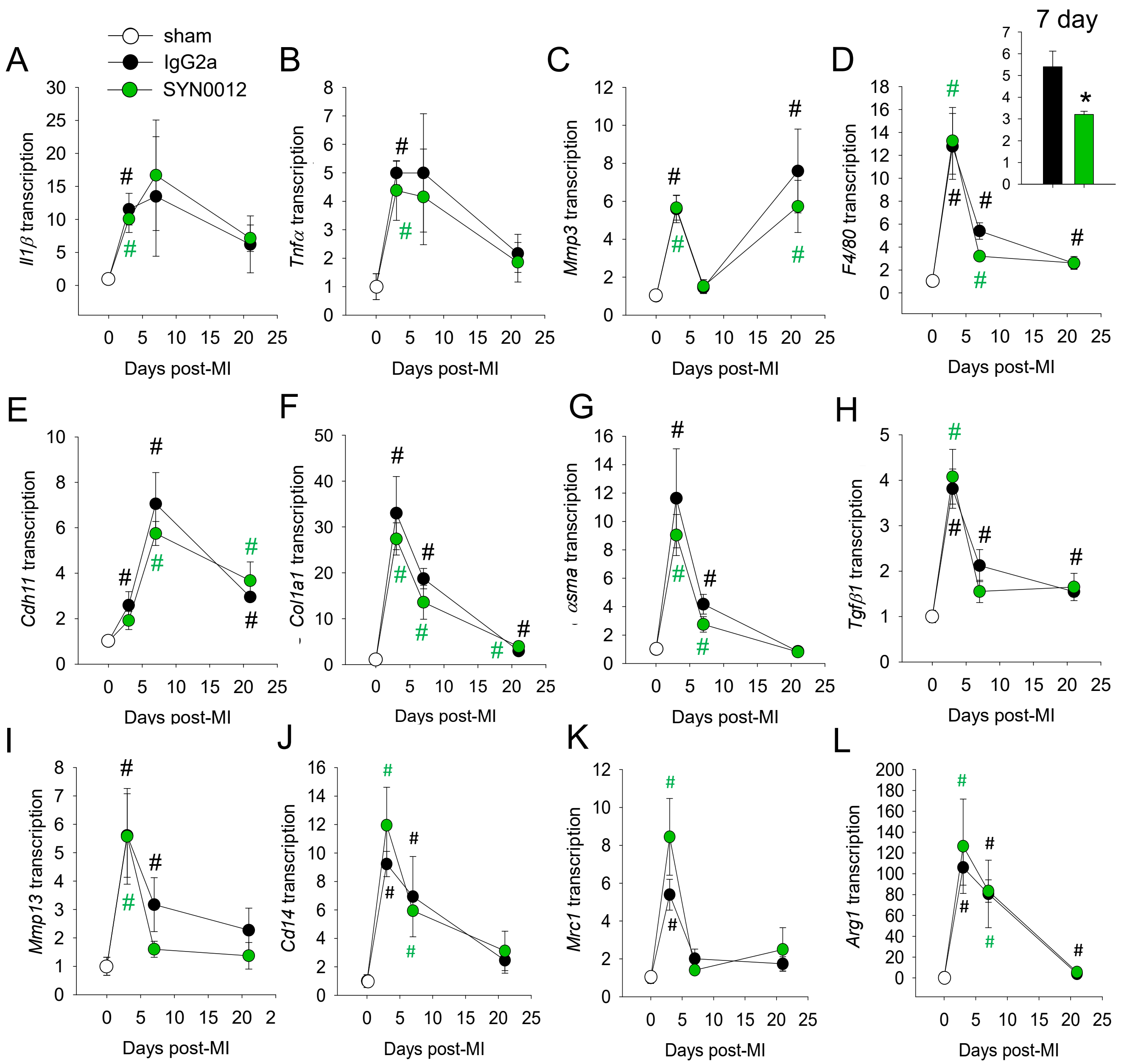


**Figure S14: Transcriptional changes of inflammatory and profibrotic genes after MI. A.** *Il1β*, **B.** *Tnfα*, **C.** *Mmp3*, **D.** *F4/80*, **E.** *Cdh11*, **F.** *Col1a1*, **G.** *αsma*, **H.** *Tgfβ1*, **I.** *Mmp13*, **J.** *Cd14*, **K.** *Mrc1*, and **L.** *Arg1*. * $p < 0.05$ between treatments, # $p < 0.05$ relative to Sham. n $\geq$ 3 for all samples.

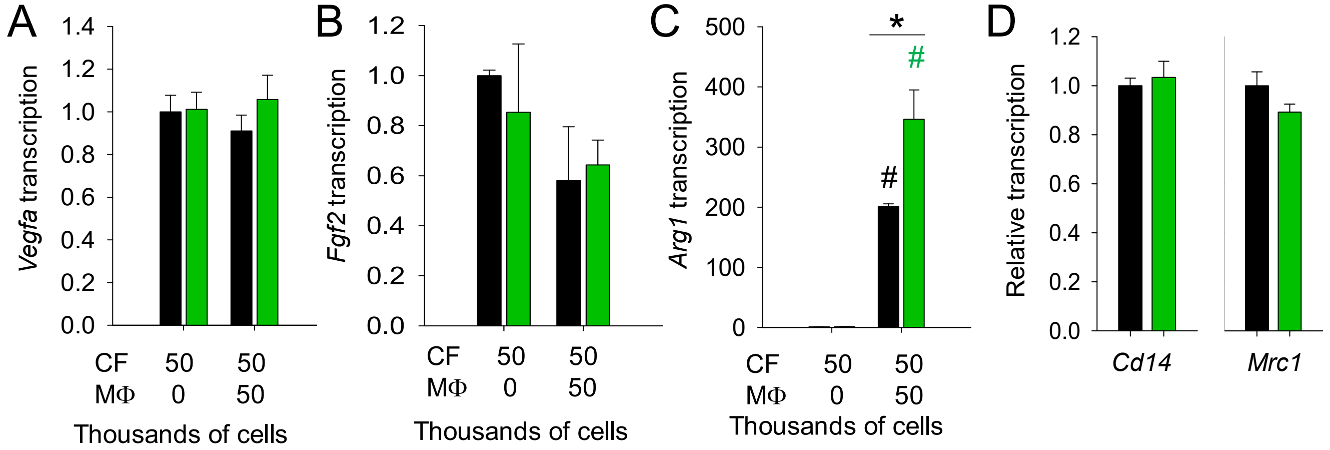


**Figure S15: Effect of CF-MΦ co-culture on transcription of proangiogenic and macrophage markers.** There were no transcriptional differences in proangiogenic growth factors *Vegfa1* (**A**) and *Fgf2* (**B**) with SYN0012 treatment. Conversely, there was a significant increase in *Arg1* transcription – a marker of M2 macrophage polarization – with SYN0012 treatment (**C**), but no difference in M1 and M2 macrophage markers *Cd14* and *Mrc1*, respectively (**D**). * $p > 0.05$ between treatments, # $p > 0.05$ relative to CF-only group; color of marker denotes treatment group. n = 3 for all samples.
